# Mapping Contributions of the Anterior Temporal Semantic Hub to the Processing of Abstract and Concrete Verbs

**DOI:** 10.1101/2024.09.02.610833

**Authors:** Emiko J. Muraki, Penny M. Pexman, Richard J. Binney

**Author notes:** Correspondence concerning this article should be addressed to Emiko J. Muraki, University of Calgary, 2500 University Drive, Calgary, Alberta, Canada, T2N 1N4 and Richard J. Binney, School of Psychology and Sport Science Bangor University Gwynedd, LL57 2AS, Wales, UK.

## Abstract

Multiple representation theories of semantic processing propose that word meaning is supported by simulated sensorimotor experience in modality-specific neural regions, as well as in cognitive systems that involve processing of linguistic, emotional, and introspective information. According to the Hub and Spoke Model of Semantic Memory, activity from these distributed cortical areas feeds into a primary semantic hub located in the ventral anterior temporal lobe (vATL). Though a substantial amount of research has tested this model in terms of concrete noun representation, there is less known about how this model can account for the representation of verb meaning, and in particular the meaning of abstract verbs which convey important information, for example, about socioemotional dynamics. In the present pre-registered study, we examined whether different types of abstract verbs (mental, emotional, nonembodied) and concrete (embodied) verbs all engage the vATL, and also whether they differentially recruit a broader set of distributed neurocognitive systems (consistent with multiple representation theories). Finally, we investigated whether there is information about different verb types distributed across the broader ATL region, consistent with a Graded Semantic Hub Hypothesis. We collected data from 30 participants who completed a syntactic classification task (is it a verb? Yes or no) and a numerical judgement task which served as an active but less semantic baseline task. Whole brain univariate analyses revealed consistent BOLD signal throughout the canonical semantic network, including the left inferior frontal gyrus, left middle temporal gyrus, and the vATL. All types of abstract verbs engaged the vATL except for mental state verbs. Finally, a multivariate pattern analysis revealed clusters within the ATL that were differentially engaged when processing each type of abstract verb. Our findings extend previous research and suggest that the hub-and-spoke hypothesis and the graded semantic hub hypothesis provide a neurobiologically constrained model of semantics that can account for abstract verb representation and processing.

---

Language plays a major role in how we interact with the world around us, yet we lack a complete cognitive and neural explanation for how we represent the meaning of words and phrases. A key unanswered question is how we understand abstract words, that is, words that cannot be directly experienced through the senses (e.g., *idea* or *regret*). Strongly embodied theories of semantic processing propose that word meaning is instantiated in simulations of prior sensory and motor experience (Barsalou et al., 2003; Glenberg & Gallese, 2012). While these theories are well suited to accounting for representation of concrete concepts (e.g., *apple* or *kick*), they cannot be readily extended to abstract concepts. In contrast, amodal theories posit that conceptual representations have a qualitatively distinct format and are divorced from sensory and motor information. They can account for abstract concepts because semantic representations can be composed only of linguistic associations, as one example. Importantly, they predict a lack of involvement of modality-specific systems during language processing, which is inconsistent with current neuroimaging and neuropsychological evidence (see below; Meteyard et al., 2012; Muraki et al., 2023). Multidimensional theories, on the other hand, propose that word meaning is grounded in sensory, motor, emotional, and other introspective experiences, as well as linguistic representations (for an overview see Meteyard et al., 2012; Muraki et al., 2023). This occurs across a distributed semantic network that includes modality-specific and multimodal associative brain regions (Binder & Desai, 2011; Tong et al., 2022), and these theories are thereby able to account for both concrete and abstract concepts. Some also make predictions about their brain basis. One highly influential example is the hub-and-spoke account, which offers a neurobiologically constrained multidimensional model of semantic memory, supported by a wealth of multi-method evidence (Lambon Ralph et al., 2017). According to this theory, semantic processing involves (i) a distributed set of modality-specific brain regions (i.e., the spokes) and (ii) a supramodal semantic hub which is located in the bilateral anterior temporal lobes (ATL; Binney et al., 2010; Lambon Ralph et al., 2017; Patterson et al., 2007; Patterson & Lambon Ralph, 2016). The purpose of the present study is to examine 1) if the ATL is engaged as a semantic hub for processing abstract words and 2) whether different abstract word types differentially engage the semantic network, both at the level of cortical and subcortical ‘spoke’ regions, as well as within the ATL hub.

## The ATL Semantic Hub

A defining feature of the hub-and-spoke account is the proposal that the bilateral ATLs play a centrally important role as a supramodal hub for semantic processing. The ATL hub serves two main functions: 1) it mediates transmodal interactions between different types of input from the ‘spokes’, and 2) it integrates multimodal information across time and contexts resulting in the distillation of coherent, generalizable concepts (Lambon Ralph, 2014; Lambon Ralph et al., 2010). These functions may be particularly important for abstract concepts whose meaning is more heavily dependent on linguistic and socioemotional experience and emerges from exposure over diverse contexts (Dove et al., 2022; Hoffman et al., 2013; Pexman et al., 2023). There is now considerable evidence for the role of the ATL in processing concrete nouns, but its role in processing other types of concept, like action concepts (i.e., verbs) and abstract concepts, is less established (Hoffman et al., 2015).

The longest standing evidence that the ATL serves as a hub for semantic processing comes from cognitive neuropsychology and the detailed study of patients with semantic dementia (SD). SD is a neurodegenerative disorder which selectively impairs semantic memory, and affects concepts of all kinds (Jefferies et al., 2009). As a consequence, it presents with profound language and communication deficits that manifest in both production and comprehension (Hodges et al., 1992; Warrington, 1975). The disorder is coupled with focal atrophy and hypometabolism of the bilateral ATL (Patterson et al., 2007). Until recently, however, there was a paucity of convergent evidence from functional neuroimaging regarding the ATL hub. This is largely due to severe and localised signal loss and image distortion that occurs with standard fMRI acquisitions (Devlin et al., 2000). However, the development of ATL-optimized fMRI techniques has paved the way for a series of neuroimaging studies that demonstrate pervasive ATL activity in response to a range of meaning-imbued stimuli, including pictures, environmental sounds, and both concrete and abstract word stimuli (Hoffman et al., 2015; Visser, Embleton, et al., 2010; Visser & Lambon Ralph, 2011). This is further bolstered by direct intracranial recordings, surface electrophysiology, and neuromodulation studies of neurotypical adults (Binney et al., 2010, 2012, 2016; Binney & Lambon Ralph, 2015; Chen et al., 2013, 2016; Marinkovic et al., 2003; Pobric et al., 2007; Rahimi et al., 2022; Rice et al., 2018).

Methodological developments have also facilitated more detailed accounts of the topology of ATL function, including the graded semantic hub hypothesis (Bajada et al., 2017; Binney et al., 2012; Jackson et al., 2018; Patterson & Lambon Ralph, 2016; Rice et al., 2015). According to a graded hub view, a large and bilateral volume of anterior temporal cortex comprises a unified representational space, all of which is engaged by semantic processing. However, there exists a center-point of this space, located within the ventrolateral ATL (inclusive of the anterior fusiform and inferior temporal gyri), which is engaged by semantic information of any kind and is invariant to, for example, the modality through which concepts are accessed. Away from the center and towards the edges, there are gradual shifts in semantic function such that regions on the periphery are relatively more specialized (Bajada et al., 2019; Binney et al., 2012; Rice et al., 2015), and encode certain types of semantic features, such as those primarily experienced through particular sensorimotor modalities (e.g., vision or audition; for a computational exploration of this general hypothesis, see Plaut, 2002). As we shall explore in more detail in the next section, the graded hub account can account for a growing number of observations of differential engagement of ATL subregions for concepts of different types, including abstract and social concepts (Binney, 2016; Hoffman et al., 2015).

### The Nature and Neural Basis of Abstract Concepts

Historically, abstract concepts have been defined in terms of what they are not; they are not concrete, in that they cannot be experienced through the senses. Thus, alternative mechanisms for the learning and retrieval of their meaning have been proposed, including a dependence on language (Paivio, 1971). In terms of their brain basis, traditional accounts predict that semantic processing of concrete and abstract words will differentially recruit perceptual-motor and verbal neural systems. However, contemporary accounts challenge dichotomic approaches to the abstract/concrete distinction (Banks & Connell, 2023; Barsalou et al., 2018), and argue that there are (i) additional sources of information that contribute to the meanings of concepts, (ii) that concreteness is a continuum, and (iii) that there is greater heterogeneity than previously thought, such that there may be different types of abstract (and concrete) concepts (Kiefer & Harpaintner, 2020). For example, behavioural evidence has shown that both abstract and concrete concepts can be strongly associated with visual sensory experience, that abstract concepts are more strongly associated with hearing, and actions of the mouth and head (Banks & Connell, 2023), and that interoception (i.e., sensations inside the body; Connell et al., 2018) and social experience (Diveica et al., 2024a, 2024b) play a key role in understanding some concepts. There is also convergent neural evidence for these claims. For example, abstract concepts associated with motor or visual experience (e.g., *fitness* or *beauty* respectively) activate the corresponding sensorimotor systems (Harpaintner et al., 2020), and face regions of the motor cortex are activated by abstract words related to mental states or processes (e.g., *logic*; Dreyer & Pulvermuller, 2018; Trumpp et al., 2024), with the rationale that these concepts may be grounded in simulations of cognitive, social or linguistic experience, involving, for instance, facial and mouth movements. Moreover, recent systematic reviews and meta-analyses of neuroimaging studies identify a distributed network of semantic activation that extends beyond visual, auditory, and somatosensory and motor cortex. For instance, emotion concepts engage the bilateral amygdala, and in some cases the orbitofrontal cortex (Conca et al., 2021; Desai et al., 2018; Kuhnke et al., 2022). Mental state concepts are associated with activity in parts of the theory of mind network (e.g., medial prefrontal cortices and the temporo-parietal junction (Conca et al., 2021; Desai et al., 2018), and processing numerical concepts has been associated with parietal regions implicated in both spatial cognition and mental calculations (Conca et al., 2021; Desai et al., 2018). Unfortunately, these studies were restricted to noun processing (Conca et al., 2021) or included sentences and story-level stimuli (Desai et al., 2018), such that there remains little understanding of the way in which some concepts, like individual verbs, are processed (Zwaan, 2014). Nonetheless, these studies suggest, overall, that word meaning may be grounded not only in sensorimotor and linguistic experience, but also via emotional, social, and other mental experiences.

The hub-and-spoke account can accommodate multiple sources of semantic information (or ‘spokes’), but its central premise is that integration of this information takes place in the ATL hub, and this is irrespective of the nature or type of concept. Neuropsychological evidence for the role of the ATL in abstract as well as concrete concept processing is inconsistent, but also complex, as is often observed of patient data (see Hoffman, 2016; Mancano & Papagno, 2023 for a review). Neuroimaging evidence from healthy populations, however, is able to demonstrate equivalent engagement of the hub’s vATL centrepoint for both abstract and concrete concepts (Binney et al., 2016; Hoffman et al., 2015; Rice et al., 2018). Key to these studies was the use of methods optimized for detecting BOLD signal in the vATL. Other studies have shown differences in the ATL, with abstract concepts more strongly engaging the dorsolateral ATL than concrete concepts (Binder et al., 2009; Bucur & Papagno, 2021; J. Wang et al., 2010). Most fMRI studies of abstract concepts, however, have not been ATL-optimised and therefore, there is still much to be gleaned about the way different subregions are engaged by different kinds of concepts.

There is evidence that different types of abstract concept engage different ATL sub-regions, in a way that is consistent with the graded semantic hub hypothesis. Social abstract concepts have been associated with increased activity in the dorsolateral and polar ATL (Binney et al., 2016) as well as the left superior temporal sulcus (X. Wang et al., 2019). Emotional abstract concepts have been associated with the left temporal pole (Conca et al., 2021; Desai et al., 2018; Kuhnke et al., 2022; X. Wang et al., 2019), and mental state abstract concepts have been associated with various ATL subregions, including the middle and inferior temporal gyri (Desai et al., 2018). According to the graded semantic hub hypothesis, differential engagement of ATL subregions to certain types of semantic information reflects preferential connectivity to inputs from sensorimotor, linguistic and other pre-semantic processes (Binney et al., 2012). The sensitivity of dorsal/polar subregions to socioemotional features may, for example, reflect dense connectivity to frontal limbic regions (via the uncinate fasciculus; Bajada et al., 2019; Papinutto et al., 2016). On the other hand, the vATL centrepoint is equally connected to all inputs and therefore should respond equally to all kinds of concepts. It has yet to be fully and systematically explored, however, whether (i) the vATL is engaged by all the above concept types (while using appropriate ATL-optimised methods), (ii) the differential engagement reflects abstractness more generally, and (iii) similar differences arise for verbs.

### The Present Study: Abstract Verbs

Abstract verbs are an understudied part of language, which contrasts with fairly extensive (including brain-based) investigations of both abstract noun and action (concrete) verb processing (Li et al., 2022; Muraki et al., 2020). This is despite the fact that abstract verbs feature frequently in our vocabulary and can convey important information about changes in mental states and other circumstances that are not directly observable, such as socioemotional dynamics. A small number of fMRI studies have investigated specific types of abstract verbs. These have shown that abstract emotion verb processing is associated with activity in the left temporal pole and middle temporal gyrus, as well as widespread activity through the motor and somatosensory cortices when compared to baseline conditions or motion verbs (Moseley et al., 2012; Rodriguez-Ferreiro et al., 2011). This suggests that both emotional and motor experiences contribute to processing these types of abstract verbs. Mental state (or cognition) verbs have been associated with activity in the left posterolateral temporal cortex when compared to motion verbs (Grossman et al., 2002). Further, change-of-state verb processing has been associated with activity in the ventrotemporal cortex (Kemmerer et al., 2008), a region also involved with motion perception. These studies did not compare different abstract verb types, nor did they closely examine the role of the ATL hub.

More recently, Muraki et al. (2020) used electroencephalogram (EEG) activity to investigate the neural correlates of processing emotion verbs, mental state verbs, and nonembodied abstract verbs (i.e., verbs not related to experiences of the human body, including change-of-state verbs such as *dissolve* or *evaporate*). Nonembodied abstract verbs had a significantly more negative N400 amplitude compared to the mental state verbs, detected at frontal electrodes. The N400 is an event-related potential component that has been associated with integrating new information during semantic processing. Furthermore, consistent with the aforementioned fMRI studies, a distributed source analysis identified the temporal pole and inferior temporal cortex as being more associated with processing mental state abstract verbs, whereas processing nonembodied abstract verbs was associated with posterior ventrotemporal cortex. Therefore, there is preliminary evidence to suggest at least partially distinct network supporting different verb types.

In the present study, we aimed to identify neural correlates of processing different types of abstract and embodied (or concrete) verbs and, in doing so, test the predictions of multidimensional theories of semantic processing, and the graded hub-and-spoke account in particular. We extended the study of Muraki et al. (2020) using distortion-corrected fMRI to reveal the involvement of the distributed cortical and subcortical ‘spoke’ regions, and the ATL semantic hub. Our pre-registered hypotheses were as follows. If multidimensional theories are correct, then different verb types will differentially recruit a distributed set of cortical and subcortical regions (outside the anterior temporal lobe). If the hub-and-spoke account is correct, then all verb types will engage the ATL semantic hub. Third, if the graded semantic hub hypothesis is correct, the vATL will be engaged by all verb types, whereas other ATL subregions may show differential activation associated with particular types of abstract verbs.

## Materials and Method

### Preregistration and Open Practices Statement

This study (hypotheses, method, and analysis plans) was pre-registered via the Open Science Framework (available at: https://osf.io/j9kub). Our stimuli are available for download at the Open Science Framework project for the current study, as are the resulting statistical maps from our analyses (https://osf.io/vd9ba/).

### Design Considerations

The approach taken in the present study was optimized to be sensitive to Blood-Oxygen Level-Dependent (BOLD) signal across all parts of the ATL, given this brain region is especially prone to magnetic susceptibility-induced signal loss and image distortion in echo planar imaging (Devlin et al., 2000; Visser, Jefferies, et al., 2010). We used the same acquisition parameters and distortion correction procedures as Balgova et al. (2022), which obtained a good signal to noise ratio in the ATL. We used a dual-echo gradient-echo echo-planar imaging (EPI) fMRI sequence, which acquires images at both a short echo-time (12ms) that is less prone to signal loss due to spin dephasing, and a long echo-time (35ms) that is more sensitive to BOLD contrast. This dual-echo sequence is more effective at detecting signal in inferior temporal regions compared to standard single gradient-echo sequences and spin-echo sequences (Halai et al., 2014, 2015). Moreover, we acquired the images with a left-to-right phase encoding direction, which reduces signal pileup in the inferior ATL (Balgova et al., 2022; Embleton et al., 2010). Finally, we applied a post-acquisition k-space spatial correction procedure to address geometric distortions in the data, which provides a more effective method of distortion correction compared to other commonly used procedures, including those based on B0 field maps (Embleton et al., 2010).

### Participants

Thirty participants took part in the experiment (21 female, 8 male, 1 nonbinary, mean age = 22.7, SD age = 4.25). All participants were between the ages of 18 – 40, first-language English speakers, with normal or corrected-to-normal vision, no history of neurological or psychiatric conditions, and were assessed to be right-handed using the Edinburgh Handedness Inventory (Oldfield, 1971). Two participants were excluded from data analysis due to poor task performance (as per our exclusion criteria in the preregistration), leaving 28 participants remaining in the final analysis (19 female, 8 male, 1 nonbinary, mean age = 22.39, SD age = 3.46). The study was conducted at Bangor University and was approved by the local research ethics review committee. Participants received monetary compensation in exchange for study participation.

### Experimental and Baseline Tasks

Participants completed a two-alternative forced-choice syntactic classification task (SCT). On each trial they were presented with a written word in the middle of a computer screen and asked to decide if the word was a verb, to which they responded either yes or no. We did not mention nouns in our framing of the task decision, as prior research has shown that this can attenuate semantic effects related to verb processing (Muraki et al., 2022). Rather, the task was framed to focus on categorization of verb or not. The baseline task was a 2-alternative forced choice numerical judgement task (NJT). On each trial they were presented with a number in the middle of a computer screen and asked to decide if the number was odd (or alternatively, asked to decide if a number was even), to which they responded either yes or no. The NJT task decision was counterbalanced across participants, such that half the participants made an ‘odd’ judgment and half made an ‘even’ judgment. The NJT was selected as a baseline task to provide a comparable visual stimulus and decision type to that of the SCT, and with the assumption that it would require similar levels of attention and general cognitive effort, but minimal semantic processing.

## Stimuli

### Words

The word stimuli were 160 verbs and 120 nouns. This included four types of verbs: 40 embodied verbs and 40 of each mental, emotional, and nonembodied abstract verbs. The different verb types were differentiated based on ratings obtained for three lexical semantic dimensions (embodiment, cognition, and valence, which are described in greater detail below) by Muraki and Pexman (2024). Each verb was in the active, present infinitive form, and had a minimum of ten valid ratings on each dimension of interest. In addition, all word stimuli (verbs and nouns) were matched (i.e., no significant differences) on lexical semantic dimensions known to influence lexical and semantic processing: word length, word frequency (log word frequency in move subtitles; Brysbaert & New, 2009), and the age of acquisition of the word (Kuperman et al., 2012). The stimuli selection process was completed using the LexOPS package (Taylor et al., 2020) for R (Version 4.2.1; R Core Team, 2022). See descriptive statistics in Table 1 for all lexical semantic dimensions.

**Table 1.** Descriptive Statistics for Word Stimuli.

| Word Type<br>(*n* = 40 per verb type<br>*n* = 60 per noun type) | Emb. *M*<br>*SD* | Valence *M*<br>*SD* | Cog. *M*<br>*SD* | Freq *M*<br>*SD* | Length *M*<br>*SD* | AoA *M*<br>*SD* | BOI *M*<br>*SD* |
| --- | --- | --- | --- | --- | --- | --- | --- |
| Mental abstract verbs | 2.86<br>0.55 | 5.28<br>0.62 | 5.59<sup>c</sup><br>0.39 | 1.58<br>0.52 | 7.38<br>1.33 | 10.85<br>1.36 | na |
| Emotional abstract verbs | 3.12<br>0.52 | 2.94<sup>b</sup><br>0.62 | 2.65<br>0.49 | 1.43<br>0.52 | 7.20<br>1.56 | 10.68<br>1.58 | na |
| Nonembodied abstract verbs | 3.11<br>0.60 | 5.29<br>0.70 | 2.67<br>0.42 | 1.42<br>0.51 | 7.30<br>1.42 | 10.84<br>1.48 | na |
| Embodied verbs | 5.20<sup>a</sup><br>0.80 | 5.12<br>0.58 | 2.48<br>0.55 | 1.43<br>0.45 | 7.28<br>1.41 | 10.57<br>1.47 | na |
| High BOI nouns | na | na | na | 1.50<br>0.41 | 7.30<br>1.41 | 10.71<br>1.29 | 5.13<sup>d</sup><br>0.62 |
| Low BOI nouns | na | na | na | 1.52<br>0.43 | 1.45<br>7.28 | 10.71<br>1.35 | 2.27<br>0.49 |

*Note.* Ratings included are embodiment (Emb.), valence, cognition (Cog.), frequency (Freq.), Length, age of acquisition (AoA), and body-object interaction (BOI). *M* = Mean, *SD* = Standard deviation. <sup>a</sup> indicates that the embodiment ratings for the embodied verbs are significantly higher than all other verbs ( $p < .05$ ). <sup>b</sup> indicates that the valence ratings for the emotional abstract verbs are significantly lower than for all other verbs ( $p < .05$ ). <sup>c</sup> indicates that the cognitive ratings for the cognitive abstract verbs are significantly higher than for all other verbs ( $p < .05$ ). <sup>d</sup> indicates that the BOI ratings for the high BOI nouns are significantly higher than for the low BOI nouns ( $p < .05$ ). There are no significant differences between any of the stimuli groups on length, frequency, or age of acquisition.

### Embodied Verbs

Embodiment ratings were based on the definition from Sidhu et al. (2014) and capture the degree to which the meaning of a verb involves the human body, rated on a scale from 1 to 7. The present set of embodied verbs had embodiment ratings greater than 4 (i.e., they were more embodied) whereas all other verb types had embodiment ratings less than 4 (i.e., they were less embodied). The embodied verbs were significantly (*p* < .05; see Table 1) higher in embodiment ratings than were the other three verb types.

### Mental Abstract Verbs

Cognition ratings were based on the definition from Muraki et al. (2020) and capture the degree to which the meaning of the verb is related to mental actions or processes of acquiring knowledge and understanding through experience and thought. Words were rated on a scale from 1 to 7. Mental abstract verbs had cognition ratings greater than 5 (i.e., they were more cognitive) and all other verb types had cognition ratings equal to or less than 3.5 (i.e., they were less cognitive). The mental abstract verbs were significantly higher (*p* < .05; see Table 1) in cognition ratings than were the other three verb types.

### Emotional Abstract Verbs

Valence ratings were based on the definition from Warriner et al. (2013) and capture the degree to which a verb’s meaning is happy versus unhappy. Words were rated on a scale from 1 to 9, where the middle point of the scale (5) indicated that the verb had neutral valence (i.e., neither happy nor unhappy). Emotional abstract verbs had valence ratings less than 4 (i.e., they had negative valence) and all other verb types had valence ratings ranging from 4 to 6.5 (i.e., they had neutral valence). The emotional abstract verbs were significantly (*p* < .05; see Table 1) lower in valence ratings than were the other three verb types.

### Nonembodied Abstract Verbs

The nonembodied abstract verbs were selected to have low embodiment ratings (less than 4), low cognitive ratings (less than or equal to 3.5), and neutral valence ratings (ranging from 4 to 6.5). Therefore, the nonembodied verbs differed significantly (*p* < .05; see Table 1) from all the other verb types and specifically on the dimension that defined each of them.

### Nouns

Nouns were included to constitute trials requiring a “no” decision in the SJT (i.e., trials where the word was not a verb). Nouns were divided into two types that differed in their association with sensorimotor experience, in order to instil a degree of variability akin to that in the verbs. We used body-object interaction ratings as a semantic dimension analogous to embodiment ratings, because the latter are only available for verbs. Body-object interaction (BOI; Pexman et al., 2019) describes the ease with which a human body can interact with a word’s referent. Half of the nouns in the present study had high BOI ratings (greater than or equal to 4; *n* = 60) and half had low BOI ratings (less than or equal to 3; *n* = 60), and they differed significantly ( *p* < .05; see Table 1).

### Numbers

The number stimuli comprised 64 odd and 64 even numbers (total *N* = 128). The numbers were randomly generated and ranged from five to eight digits in length, with an equal quantity (*n* = 32) of each length in the odd and even number groups.

### Experimental Procedure

A PC running PsychoPy (Version 2022.2.4; Peirce et al., 2019) was used for presentation of stimuli and recording of responses. Participants completed four runs, each lasting approximately 7.5 minutes. Within a run, the SJT and NJT alternated in a block design with a 14 second rest after each number block. There were four SJT blocks containing between 15 and 24 trials (Mean trials = 17.5 per block, block duration range = 56.2 to 99.2 secs) while each of four NJT blocks contained 8 trials (block duration = 20 secs). Each trial began with a 500ms fixation cross (blue cross on number trials, black cross on word trials) presented in the middle of the screen and then a number or word was presented for 2000ms in black, lower-case font on a white background. After the 2000ms response window, word trials had a variable interstimulus interval (0-13100ms; mean ISI = 1840ms) and number trials had a 0ms interstimulus interval. Participants were instructed to respond as quickly as possible after the word or number appeared on the screen using two buttons designated as “yes” and “no” on an MR-compatible response box.

Nested within the block design, we employed a rapid event-related design for presentation of the five different word conditions. This was achieved using a pseudorandomized sequence of conditions (1 trial per sequence position) that spanned the four SJT blocks and was used in all four runs. We created four unique lists of 70 words (10 of each verb type and 30 nouns). We matched each word type (embodied, emotional, mental, nonembodied, high BOI nouns and low BOI nouns) on their corresponding semantic variable across lists (e.g., the four lists did not significantly differ on the valence ratings of the emotional abstract verbs within the list). We also matched across lists for the lexical variables length, frequency, and age of acquisition. Each list was used once across the four runs. The order in which lists of words were presented was counterbalanced across participants using a balanced Latin square design. The exact word for each trial was randomly selected from a condition-specific sub-list, such that no pair of participants would have seen the words in the same order. In a similar fashion, four unique lists of 32 numbers were created and the order in which they were presented was counterbalanced across participants. All study materials are available for download on our OSF project page https://osf.io/vd9ba/.

### Imaging Acquisition

All imaging was performed on a 3.0 T Phillips Elition MRI scanner with a 32-channel head coil and a SENSE factor of 2.5. We used a dual-echo gradient-echo EPI fMRI sequence to acquire 31 axial slices covering the whole brain in ascending sequential order with the following parameters: short echo time (TE) = 12ms and long TE = 35ms, repetition time (TR) = 2000ms, and flip angle = 85°. The functional EPIs were acquired with an in-plane reconstructed resolution of 2.5 x 2.5 mm and a slice thickness of 4 mm voxels (reconstruction matrix = 96 × 96; FOV (mm) = 240 × 240 × 124). We acquired five dummy scans prior to each run (to develop a steady state magnetisation), followed by 224 volumes per run. We acquired the four main functional runs with a single direction k-space traversal in the left–right phase-encoding direction. In addition, we acquired a short EPI ‘pre-scan’ with the participants at rest. The parameters of the pre-scan matched the main functional scans with the exception that they included interleaved dual direction k-space traversals. This provided us with 10 pairs of images with opposing direction distortions (10 left–right and 10 right–left) which were used in a k-space distortion correction procedure for addressing geometric distortion and mislocalisation due to magnetic susceptibility artifacts. We acquired a high-resolution T2-weighted scan to assess the accuracy of our distortion correction procedure, which contained 36 slices covering the whole brain with the following parameters: TE = 78ms, TR = 3410ms; flip angle = 90°. The T2-weighted scan was acquired with a reconstructed voxel size (mm) of 0.6 x 0.6 x 4 and a slice thickness of 4 mm voxels with a 1 mm slice gap (reconstruction matrix = 400 x 370; FOV (mm) = 240 x 240 x 179). In addition, we used a T1-weighted imaging sequence to acquire an anatomical image used to estimate the transformation of our functional images into MNI space. The T1 images contained 175 slices covering the whole brain and were acquired with the following parameters: TE = 18ms, TR = 3.5ms; flip angle = 8°, reconstructed voxel size (mm) = 0.94 x 0.94 x 1 and reconstruction matrix = 240 x 240. All images were acquired with a 30° axial tilt.

## Data Analysis

### Data cleaning

Two participants had number judgement task accuracy values which indicated that they had reversed the buttons used for their response (e.g., accuracy less than 10%), so their number judgment task data were recoded.

### Behavioural data

Behavioural analyses were conducted using the statistical software package R (R Core Team, 2023). Task accuracy and response time were assessed using linear mixed effect models which included random effects of participant and item and fixed effects of word or trial type. We excluded practice trials, trials where the response time was more than 3 SD away from the participant’s mean response time, and words with significantly lower than chance accuracy on the SCT. We only included correct trials in our response time models. Inferential statistics were calculated to compare performance in the SCT and NJT, as well as to compare between verb types in the SCT and odd and even numbers in the NJT.

### fMRI distortion correction and preprocessing

fMRI preprocessing and analyses were conducted using the statistical software package MATLAB 2019b (for distortion correction) and 2020a (The MathWorks Inc., 2020), SPM12, and the CoSMo MVPA toolbox for MATLAB (Oosterhof et al., 2016). A spatial remapping correction was computed separately for images acquired at the long and the short echo time using the method reported in Embleton et al. (2010), implemented via an in-house MATLAB script (available upon request), as well as SPM12’s (Statistical Parametric Mapping software; Wellcome Trust Centre for Neuroimaging, London, UK) 6-parameter rigid body registration algorithm. In the first step, each functional volume was registered to the mean of the 10 pre-scan volumes acquired at the same echo time. This step is both a required part of the distortion correction procedure and corrects the timeseries for differences in participant positioning in between functional runs and for minor motion artefacts within a run. Next, one spatial transformation matrix per echo time was calculated from the opposingly distorted pre-scan images. These transformations consisted of the remapping needed to correct geometric distortion and were then applied to each of the main functional volumes. This resulted in two motion and distortion-corrected time-series per run (224 volumes per echo per run), which were subsequently combined at each timepoint using a simple linear average of image pairs.

The following pre-processing steps were completed using SPM12 in Matlab 2020a. Slice-timing correction was performed, referenced to the middle slice. The T1-weighted anatomical image was co-registered to a mean of the distortion- and motion-corrected images using a six-parameter rigid-body transform and the normalized mutual information objective function. SPM12’s unified segmentation and normalization function was used to estimate a spatial transform to register the anatomical image to Montreal Neurological Institute (MNI) standard stereotaxic space and this transform was subsequently applied to the co-registered functional volumes which were then resampled to a 2 x 2 x 2 mm voxel size. As a final step, the normalized functional images were smoothed using an 8mm full-width half-maximum Gaussian filter.

### Univariate analyses

Functional data were analysed using the general linear model approach (GLM). We conducted within-subjects fixed effects analyses with all functional runs incorporated into a single GLM per participant. Onsets for each of the verb types, nouns, and number blocks were modelled and convolved with the canonical hemodynamic response function, and a high-pass filter (cutoff of 128s) was applied. The extracted motion parameters were also entered into the GLM model as regressors of no interest. We restricted the analyses to grey matter using an explicit mask generated from a group-level probabilistic tissue segment arising out of the SPM12 unified segmentation and normalization procedure, which was binarized with a 0.4 threshold.

We tested two hypotheses: 1) that different verb types will differentially recruit a distributed set of cortical regions outside the anterior temporal lobe consistent with multidimensional theories of semantic representation, and 2) that all verb types will recruit the vATL consistent with the Hub and Spoke Model of Semantic Memory. We conducted whole brain analyses and selected a priori regions of interest based on other studies that have examined the neural correlates of abstract word meaning. At the group-level analysis, we examined whole brain contrasts between each verb type relative to the active (NJT) baseline, as well as each of the abstract verb types relative to the embodied verbs, to examine whether there were regions specific to processing abstract verb meaning above and beyond processing verb meaning generally. Whole-brain multi-subject random effects analyses were conducted on each of the following contrasts of interest: embodied verbs > numbers, mental verbs > numbers, emotional verbs > numbers, nonembodied verbs > numbers, mental verbs > embodied verbs, emotional verbs > embodied verbs, and nonembodied verbs > embodied verbs. We performed one-sample t-tests on all these sets of contrast images, restricting the statistical maps to cerebral tissue using the same explicit group-level mask as used in the single subject analyses. Cluster-wise statistical significance was assessed using a cluster-defining voxel-height threshold of *p* < .001 uncorrected, and a family-wise error (FWE) corrected cluster extent threshold at *p* < .05. The resulting thresholded maps were overlaid on a MNI152 template brain in MRIcroGL for all figures (Rorden & Brett, 2000). We used the label4MRI package implemented in R (Chuang, 2024) to inform the labelling of each cluster based on its peak coordinates.

To complement the whole-brain analyses, we used a priori regions of interest (ROI) to extract and quantify the magnitude of activation within different cortical and ATL subregions, and to increase sensitivity to the hypothesized effects. The ROI analysis was implemented using the SPM MarsBar toolbox (Brett et al., 2002). We defined 7 ROIs based on coordinates extracted from published meta-analyses or fMRI studies of semantic processing. Coordinates were used to define the centre of mass of spherical ROIs with radii of 8mm (Volume = 2056 mm3). One of these ROIs corresponds to the centre-point of the graded ATL semantic hub, located in the ventral ATL, and was based on the findings of (Binney et al., 2010). The other six spherical ROIs correspond to brain regions that might, based on prior findings, be differentially recruited by semantic processing of different verb types, as detailed in the following paragraph (also see Table 2).

**Table 2.** Spherical Regions of Interest (ROI) by Contrast.

| ROI Location | Contrast | Centre of Mass (MNI) |  |  |
| --- | --- | --- | --- | --- |
|  |  | x | y | z |
| Left anterior supramarginal gyrus | Embodied > Nonembodied | -60 | -30 | 30 |
| Left posterior superior temporal sulcus | Embodied > Nonembodied | -62 | -56 | 8 |
| Left amygdala | Emotional > Embodied | -26 | -6 | -22 |
| Right amygdala | Emotional > Embodied | 24 | -4 | -28 |
| Left temporal parietal junction | Mental > Embodied | -50 | -58 | 20 |
| Left middle temporal gyrus | Nonembodied > Embodied | -61 | -44 | -2 |
| Ventral anterior temporal lobe | Embodied > Number | -36 | -15 | -30 |
|  | Emotional > Number |  |  |  |
|  | Mental > Number |  |  |  |
|  | Nonembodied > Number |  |  |  |

For the embodied verbs we selected regions associated with conceptual processing related to action and motion from the Kuhnke et al. (2022) meta-analysis that examined modality-specific brain regions that were uniquely activated by processing particular types of concepts. We selected ROIs with the highest specificity for action and motion concepts in the contrast analyses assessing modality specificity. For action concepts this was a region in the left anterior supramarginal gyrus and for motion this was an area in the left posterior superior temporal sulcus. For the emotional verbs we selected regions associated with conceptual processing related to emotion from the Kuhnke et al. (2022) meta-analysis. We selected ROIs with the highest specificity for emotional concepts in the contrast analyses assessing modality specificity. This was located outside the anterior temporal cortex, in the bilateral amygdalae. For the mental verbs we selected a region of the temporal parietal junction associated with processing theory-of-mind-related concepts by the Desai et al. (2018) meta-analysis. We anticipated that the nonembodied verbs would be the most abstract, as they have no relation to physical or interoceptive experiences of the body, so we selected a region associated with processing abstract concepts in the Bucur and Papagno (2021) meta-analysis. The middle temporal gyrus has also been associated with language comprehension in previous research (see Dronkers et al., 2004), which is consistent with the proposal that abstract concepts that lack sensory or motor experiences may rely more on language to ground their meaning.

### Multivariate analyses

Finally, we used a constrained searchlight multivoxel pattern analysis (MVPA) to investigate whether patterns of activity within the ATL could be used to classify the different verb types. The analysis used a take-one-run-out cross-validation and a linear discriminant analysis (LDA) classifier implemented via the CoSMo MVPA toolbox (Oosterhof et al., 2016). For each participant, a GLM was estimated on unsmoothed image data and β weights were extracted. The β weights were then submitted to a searchlight method MVPA restricted to the ATL, using a mask previously described by Hung et al. (2020). This mask defined a volume of interest that covered the anterior portion of the temporal lobe, including the temporal pole, anterior superior temporal gyrus, anterior middle temporal gyrus, the anterior inferior temporal gyrus, the anterior temporal fusiform cortex, and the anterior parahippocampal gyrus. The posterior boundary of the ATL was approximately MNI coordinate y = −24 along the ventral surface and y = −14 on the lateral surface and the volume was 9706 voxels (voxel size = 2×2×2mm^3^). The searchlight was defined with a spherical neighbourhood comprising the 100 voxels nearest to the centre voxel. We assessed z-transformed classification accuracy results between mental to embodied verbs, emotional to embodied verbs, and nonembodied to embodied verbs.

## Results

### Behavioural Analyses

The descriptive statistics for both the SCT and the NJT are presented in Table 3. For two verbs response accuracy was significantly below chance and so these verbs were removed from the analyses. In the SCT there were no significant effects of verb type on response time or accuracy (as referenced against embodied verbs; see Table 4 for both models). To follow-up, we performed pairwise comparisons with a Tukey correction. There were no significant differences in response time for any pairwise comparison. There was one significant difference in accuracy: Emotional abstract verbs were responded to less accurately than mental abstract verbs (*z* = -2.92, adjusted *p* = .019). Responses to the SCT were significantly slower and less accurate than the NJT (see Table 5 for both models). In the NJT odd trials were significantly more accurate than even trials, however there was no difference between odd and even numbers on response time (see Table 6 for both models).

**Table 3.** Response Time and Accuracy Descriptive Statistics by Task and Trial Type.

| Task and Trial Type | *M* Participant RT in milliseconds<br>(*SD*) | *M* Participant Accuracy in %<br>(*SD*) |
| --- | --- | --- |
| *Syntactic Classification Task* |  |  |
| All Trials | 924.40<br>(133.49) | 90.48<br>(8.44) |
| Embodied Verbs | 906.44<br>(158.21) | 90.63<br>(12.69) |
| Emotional Verbs | 920.86<br>(148.85) | 87.82<br>(14.63) |
| Mental Verbs | 898.02<br>(160.66) | 92.12<br>(13.02) |
| Nonembodied Verbs | 920.66<br>(156.63) | 88.93<br>(14.20) |
| Nouns | 948.24<br>(125.47) | 91.28<br>(7.33) |
| *Number Judgement Task* |  |  |
| All Trials | 830.90<br>(198.09) | 93.22<br>(8.02) |
| Even | 841.03<br>(208.90) | 89.90<br>(14.04) |
| Odd | 820.77<br>(192.00) | 96.54<br>(3.79) |

**Table 4.** Mixed Effects Models Predicting SCT Response Times and Accuracy.

|  | Linear Regression - Response Times |  |  | Logistic Regression - Accuracy |  |  |
| --- | --- | --- | --- | --- | --- | --- |
| Fixed Effects | *b* | 95% *CI* | *p* | *OR* | 95% *CI* | *p* |
| Intercept | 900.66 | [838.70, 962.63] | < .001 | 21.20 | [11.86, 37.88] | < .001 |
| Verb Type (Emotional) | 13.94 | [-17.69, 45.58] | .388 | 0.67 | [0.45, 1.01] | .056 |
| Verb Type (Mental) | -9.23 | [-40.70, 22.23] | .565 | 1.25 | [0.82, 1.92] | .298 |
| Verb Type (Nonembodied) | 9.72 | [-21.68, 41.12] | .544 | 0.79 | [0.52, 1.18] | .249 |

| Random Effects | Variance | SD | Variance | SD |
| --- | --- | --- | --- | --- |
| Participant Intercept | 24388 | 156.17 | 0.36 | 0.60 |
| Item Intercept | 3168 | 56.28 | 1.71 | 1.31 |
| Residual | 48484 | 220.19 |  |  |
| Model Fit | Marginal | Conditional | Marginal | Conditional |
| R<sup>2</sup> | .001 | .363 | .010 | .393 |

Note. CI = confidence interval; OR = odds ratio; Verb type is a factor variable with embodied verbs set as the reference group. Accuracy is a binary dependent variable with inaccurate responses as the reference group (0) and accurate responses as the focus group (1). The marginal R<sup>2</sup> includes only the variance from the fixed effects and the conditional R<sup>2</sup> includes variance from both the fixed and random effects. The model equations for response time and accuracy respectively were: RT ~ Verb Type + (1|Participant) + (1|Item) and Acc ~ Verb Type + (1|Participant) + (1|Item). *p*-values for fixed effects in the linear mixed effects model are calculated using Satterthwaite's method. *N* participants = 28, *N* stimuli = 158.

**Table 5.**
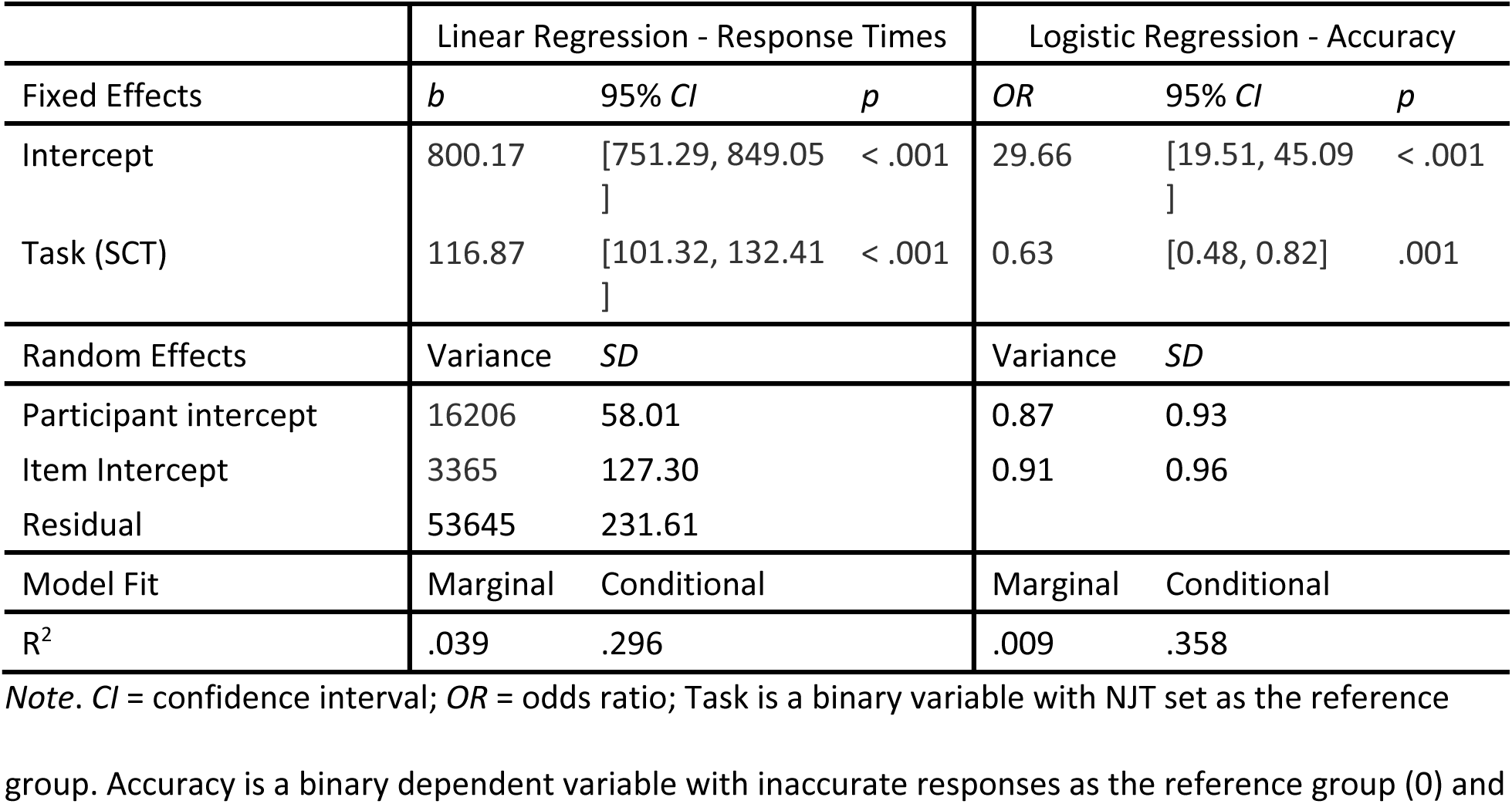
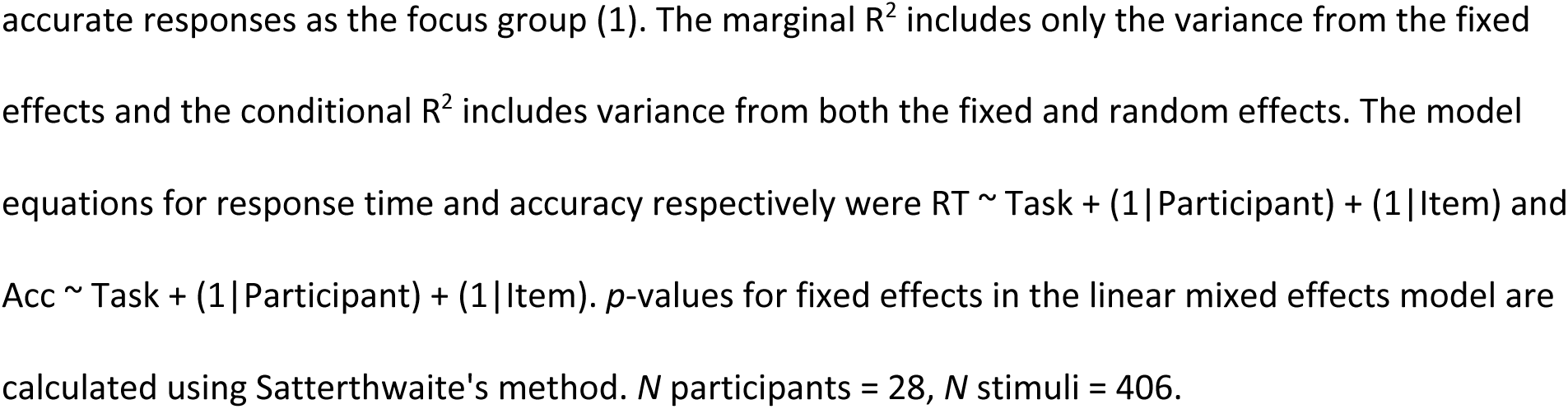
Mixed Effects Models Predicting Task Response Times and Accuracy.

**Table 6.** Mixed Effects Models Predicting NJT Response Times and Accuracy.

|  | Linear Regression - Response Times |  |  | Logistic Regression - Accuracy |  |  |
| --- | --- | --- | --- | --- | --- | --- |
| Fixed Effects | *b* | 95% *CI* | *p* | *OR* | 95% *CI* | *p* |
| Intercept | 815.04 | [746.53, 883.55] | < .001 | 19.66 | [11.03, 35.03] | < .001 |
| Number Type (Odd) | -10.17 | [-24.91, 4.58] | .177 | 3.50 | [2.43, 5.05] | < .001 |
| Random Effects | Variance | *SD* |  | Variance | *SD* |  |
| Participant intercept | 22246 | 182.61 |  | 0.28 | 0.52 |  |
| Item Intercept | 24 | 4.85 |  | 1.83 | 1.35 |  |
| Residual | 45067 | 212.29 |  |  |  |  |
| Model Fit | Marginal | Conditional |  | Marginal | Conditional |  |
| $R^2$ | .000 | .426 | | .068 | .432 | |

*Note.* *CI* = confidence interval; *OR* = odds ratio; Number Type is a binary variable with even set as the reference group. Accuracy is a binary dependent variable with inaccurate responses as the reference group (0) and accurate responses as the focus group (1). The marginal $R^2$ includes only the variance from the fixed effects and the conditional $R^2$ includes variance from both the fixed and random effects. The model equations for response time and accuracy respectively were $RT \sim \text{Number Type} + (1|\text{Participant}) + (1|\text{Item})$ and $\text{Acc} \sim \text{Number Type} + (1|\text{Participant}) + (1|\text{Item})$ . $p$ -values for fixed effects in the linear mixed effects model are calculated using Satterthwaite's method. $N$ participants = 28, $N$ stimuli = 128.

## Univariate fMRI Analyses

### Whole Brain Contrasts

Whole brain univariate analyses were conducted to contrast (i) the syntactic classification task with the active baseline (number judgement), (ii) each verb type with the active baseline and (iii) each abstract verb type with embodied verbs. Compared to the baseline task, the SCT was associated with robust activation in the semantic network (see Figure 1 and Table 7). This included a large cluster that extended up from the vATL (x ≈ -30) up into the left ventrolateral prefrontal cortex encompassing regions of the pars orbitalis and the pars triangularis. There were also clusters of activation extending along the anterior superior temporal gyrus, posterior middle, and inferior temporal gyri. In addition, we observed activation in the left occipital lobe, possibly due to the greater visual complexity of word compared to number stimuli (Binney et al., 2016) as well as activation in the right cerebellum.

**Figure 1.**
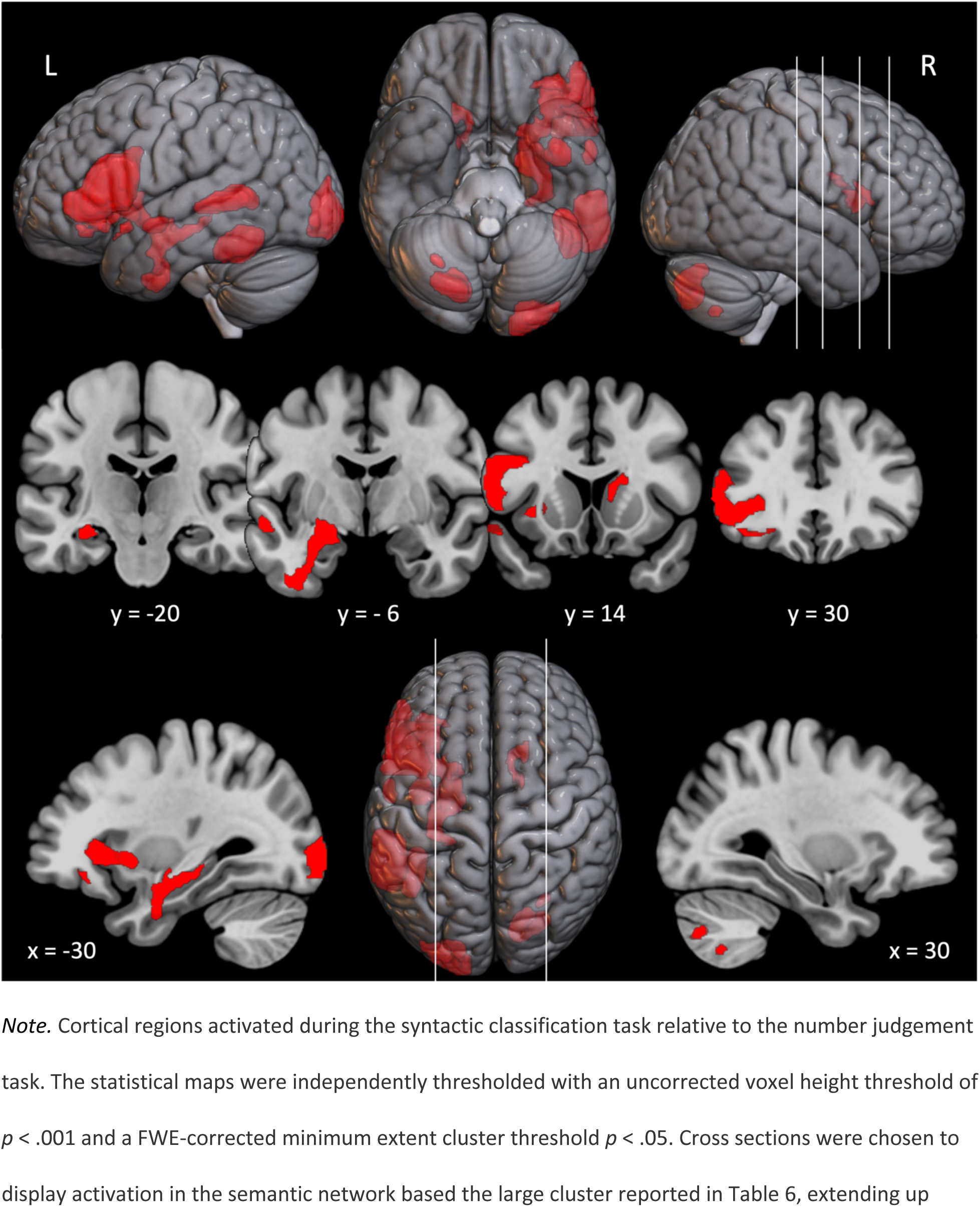
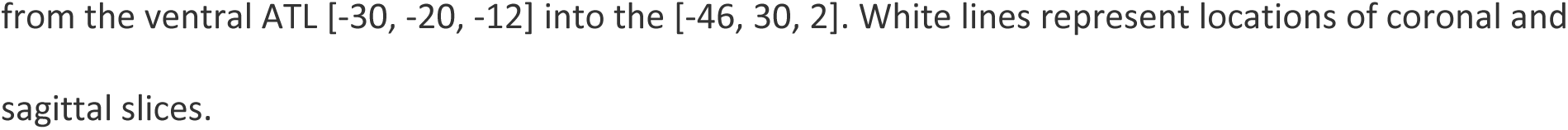
Cortical Regions Activated by Syntactic Classification > Number Judgement.

**Table 7.**
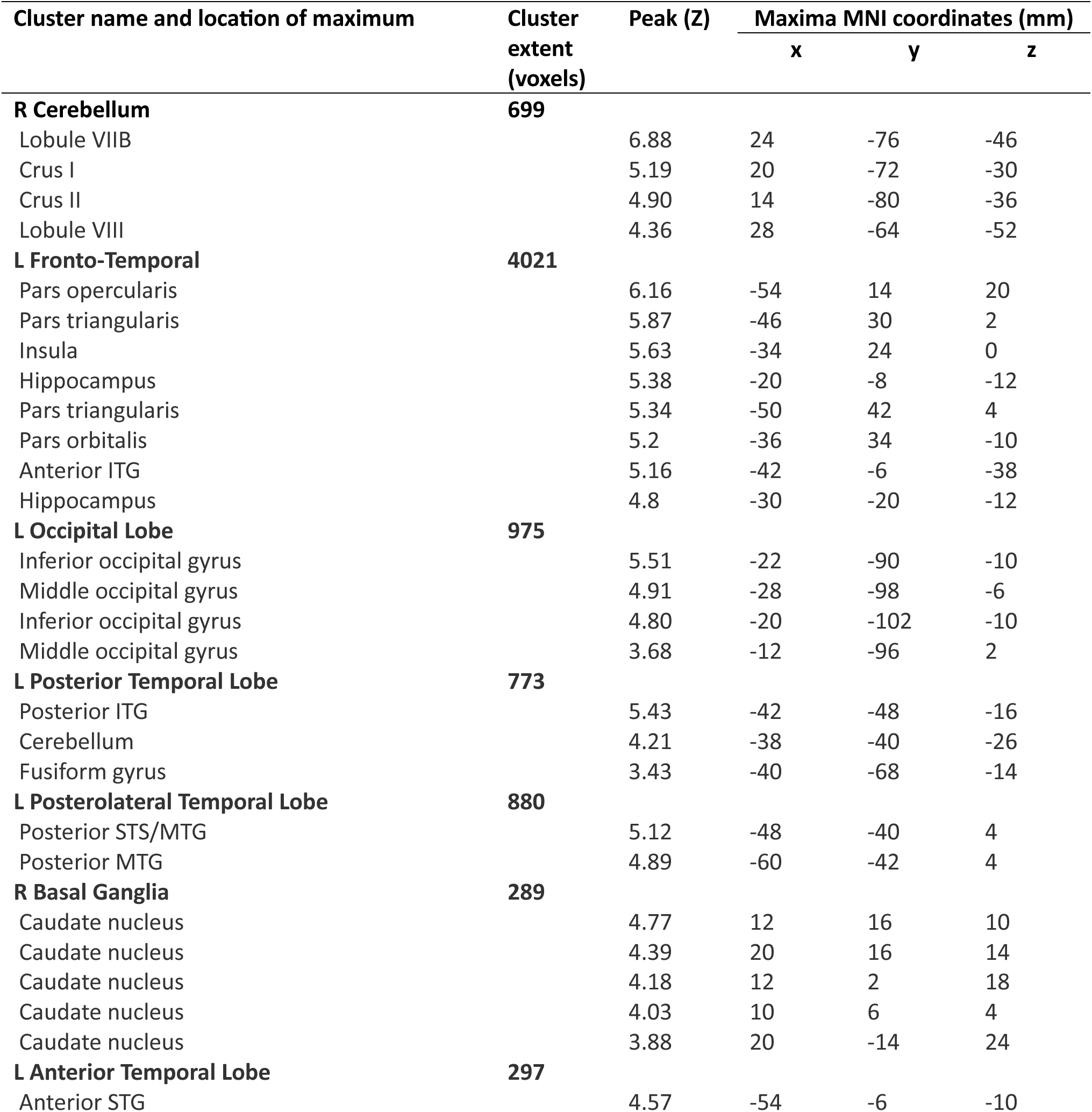
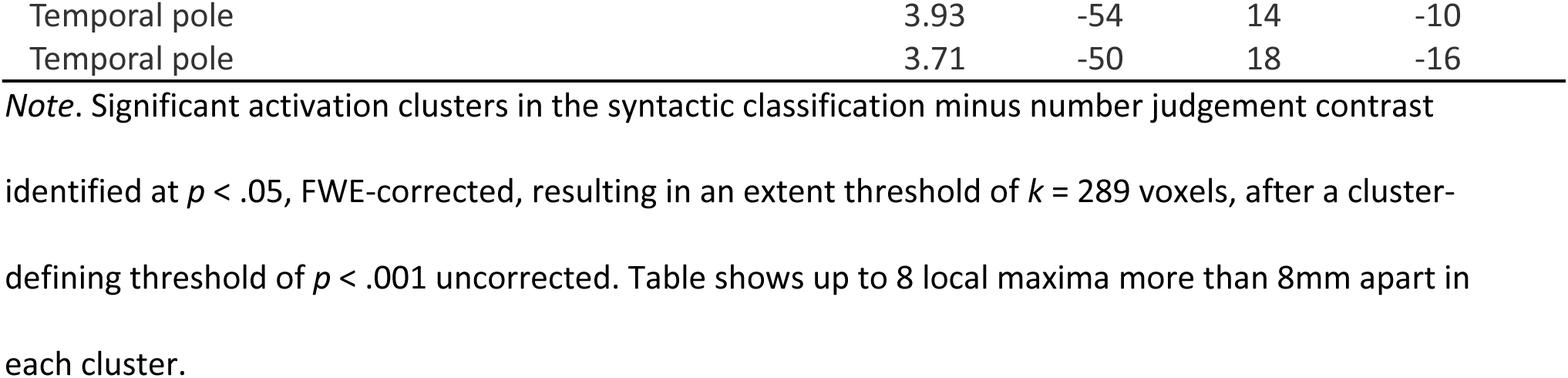
Significant Activation Clusters for Syntactic Classification Relative to Number Judgement.

| Cluster name and location of maximum | Cluster extent (voxels) | Peak (Z) | Maxima MNI coordinates (mm) |  |  |
| --- | --- | --- | --- | --- | --- |
|  |  |  | x | y | z |
| **R Cerebellum** | **699** |  |  |  |  |
| Lobule VIIb |  | 6.88 | 24 | -76 | -46 |
| Crus I |  | 5.19 | 20 | -72 | -30 |
| Crus II |  | 4.90 | 14 | -80 | -36 |
| Lobule VIII |  | 4.36 | 28 | -64 | -52 |
| **L Fronto-Temporal** | **4021** |  |  |  |  |
| Pars opercularis |  | 6.16 | -54 | 14 | 20 |
| Pars triangularis |  | 5.87 | -46 | 30 | 2 |
| Insula |  | 5.63 | -34 | 24 | 0 |
| Hippocampus |  | 5.38 | -20 | -8 | -12 |
| Pars triangularis |  | 5.34 | -50 | 42 | 4 |
| Pars orbitalis |  | 5.2 | -36 | 34 | -10 |
| Anterior ITG |  | 5.16 | -42 | -6 | -38 |
| Hippocampus |  | 4.8 | -30 | -20 | -12 |
| **L Occipital Lobe** | **975** |  |  |  |  |
| Inferior occipital gyrus |  | 5.51 | -22 | -90 | -10 |
| Middle occipital gyrus |  | 4.91 | -28 | -98 | -6 |
| Inferior occipital gyrus |  | 4.80 | -20 | -102 | -10 |
| Middle occipital gyrus |  | 3.68 | -12 | -96 | 2 |
| **L Posterior Temporal Lobe** | **773** |  |  |  |  |
| Posterior ITG |  | 5.43 | -42 | -48 | -16 |
| Cerebellum |  | 4.21 | -38 | -40 | -26 |
| Fusiform gyrus |  | 3.43 | -40 | -68 | -14 |
| **L Posterolateral Temporal Lobe** | **880** |  |  |  |  |
| Posterior STS/MTG |  | 5.12 | -48 | -40 | 4 |
| Posterior MTG |  | 4.89 | -60 | -42 | 4 |
| **R Basal Ganglia** | **289** |  |  |  |  |
| Caudate nucleus |  | 4.77 | 12 | 16 | 10 |
| Caudate nucleus |  | 4.39 | 20 | 16 | 14 |
| Caudate nucleus |  | 4.18 | 12 | 2 | 18 |
| Caudate nucleus |  | 4.03 | 10 | 6 | 4 |
| Caudate nucleus |  | 3.88 | 20 | -14 | 24 |
| **L Anterior Temporal Lobe** | **297** |  |  |  |  |
| Anterior STG |  | 4.57 | -54 | -6 | -10 |

|  |  |  |  |  |
| --- | --- | --- | --- | --- |
| Temporal pole | 3.93 | -54 | 14 | -10 |
| Temporal pole | 3.71 | -50 | 18 | -16 |

*Note.* Significant activation clusters in the syntactic classification minus number judgement contrast
identified at $p < .05$ , FWE-corrected, resulting in an extent threshold of $k = 289$ voxels, after a cluster-defining threshold of $p < .001$ uncorrected. Table shows up to 8 local maxima more than 8mm apart in each cluster.

The contrasts comparing each verb type to the baseline task show a similar pattern of activation to the contrast comparing all words with the baseline task, with a few exceptions (see Figure 2 and Table 8). All verbs were associated with activity in the inferior frontal gyrus (IFG) and posterior middle temporal gyrus (pMTG), both core regions in the semantic network. Emotional, embodied, and nonembodied verbs were associated with activity in the vATL, yet mental verbs were not. Embodied verbs were associated with a larger cluster of activation in the left IFG and with activation extending more posterior on the MTG than any other verb condition. Similarly, nonembodied verbs were associated with larger clusters of activation in the left IFG and the posterior portion of the left MTG and fusiform gyri. They were also uniquely associated with activity in the left motor cortex and the left and right somatosensory cortices. There were no significant differences in activation between each abstract verb type compared to the embodied verbs after corrections, suggesting that they showed a similar pattern of activation. However, there was one cluster in the left motor cortex that was more active for nonembodied compared to embodied verbs that approached significance (*p* = .069 FWE-corrected, resulting in an extent threshold of *k* = 133 voxels.)

**Figure 2.**
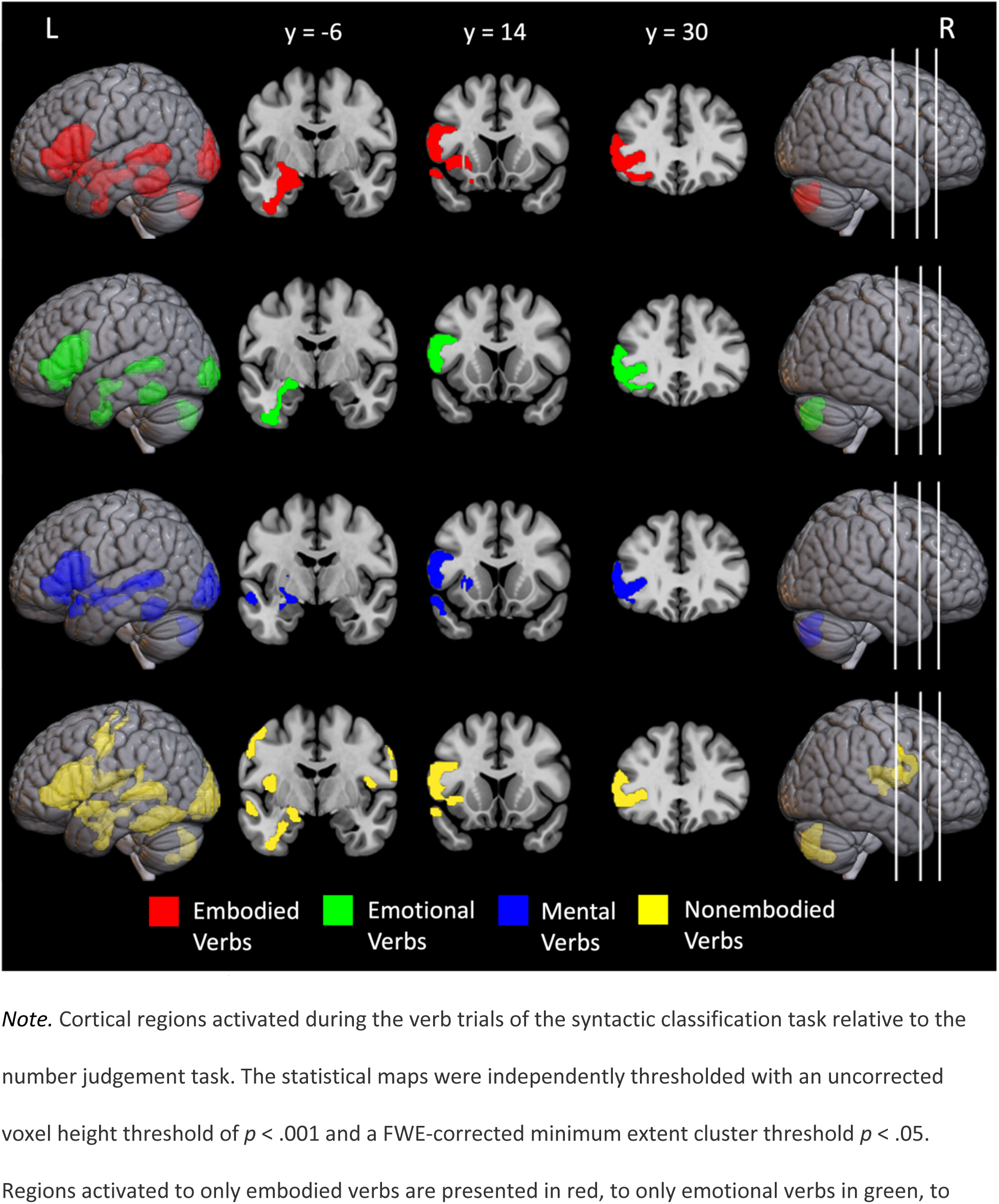
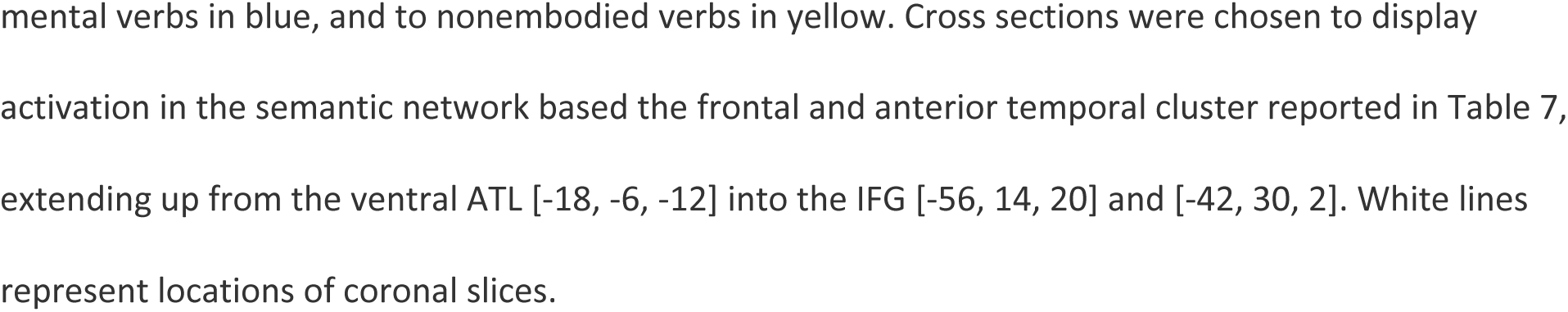
Cortical Regions Activated by Each Verb Type > Number Judgement.

**Table 8.**
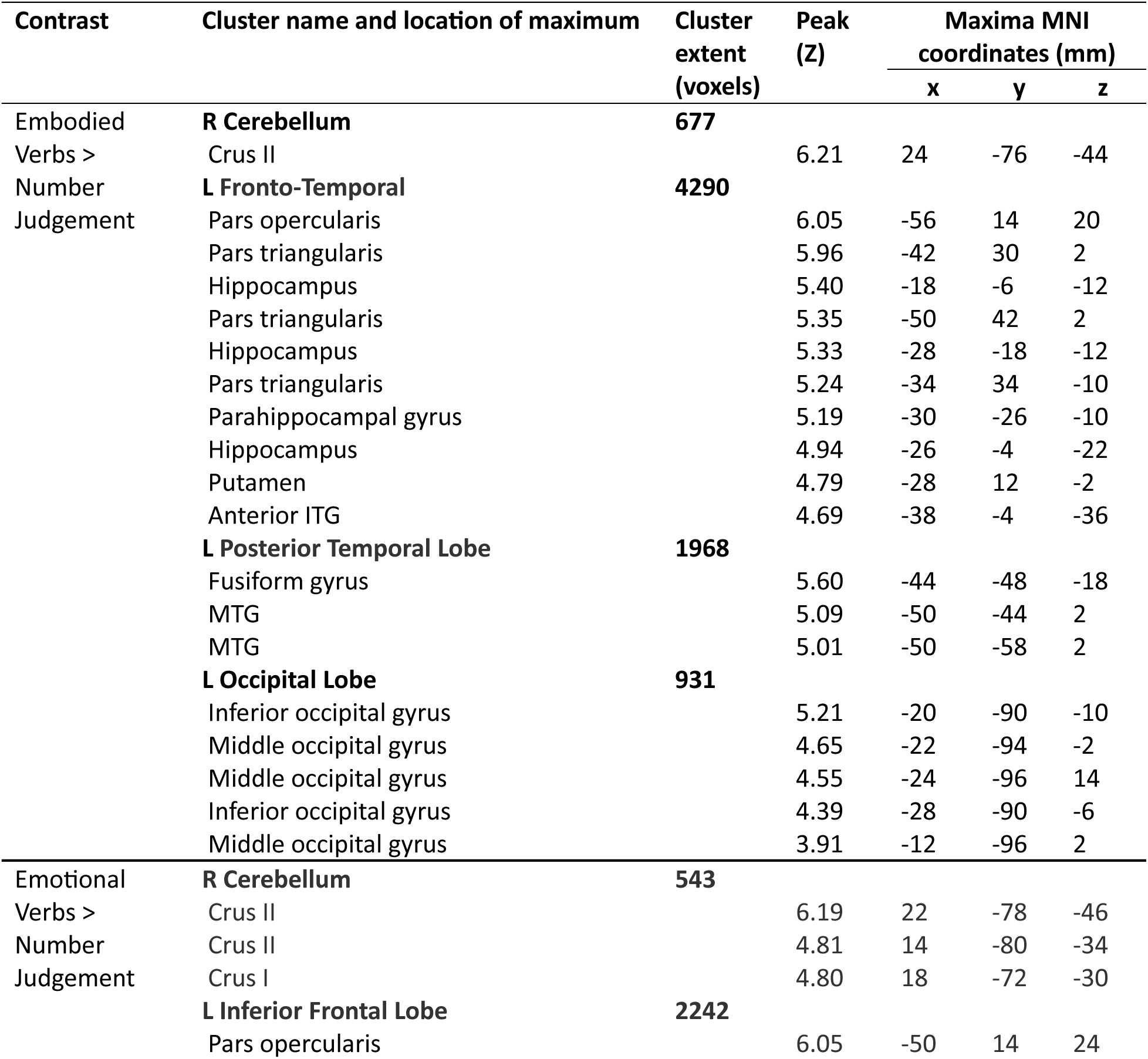
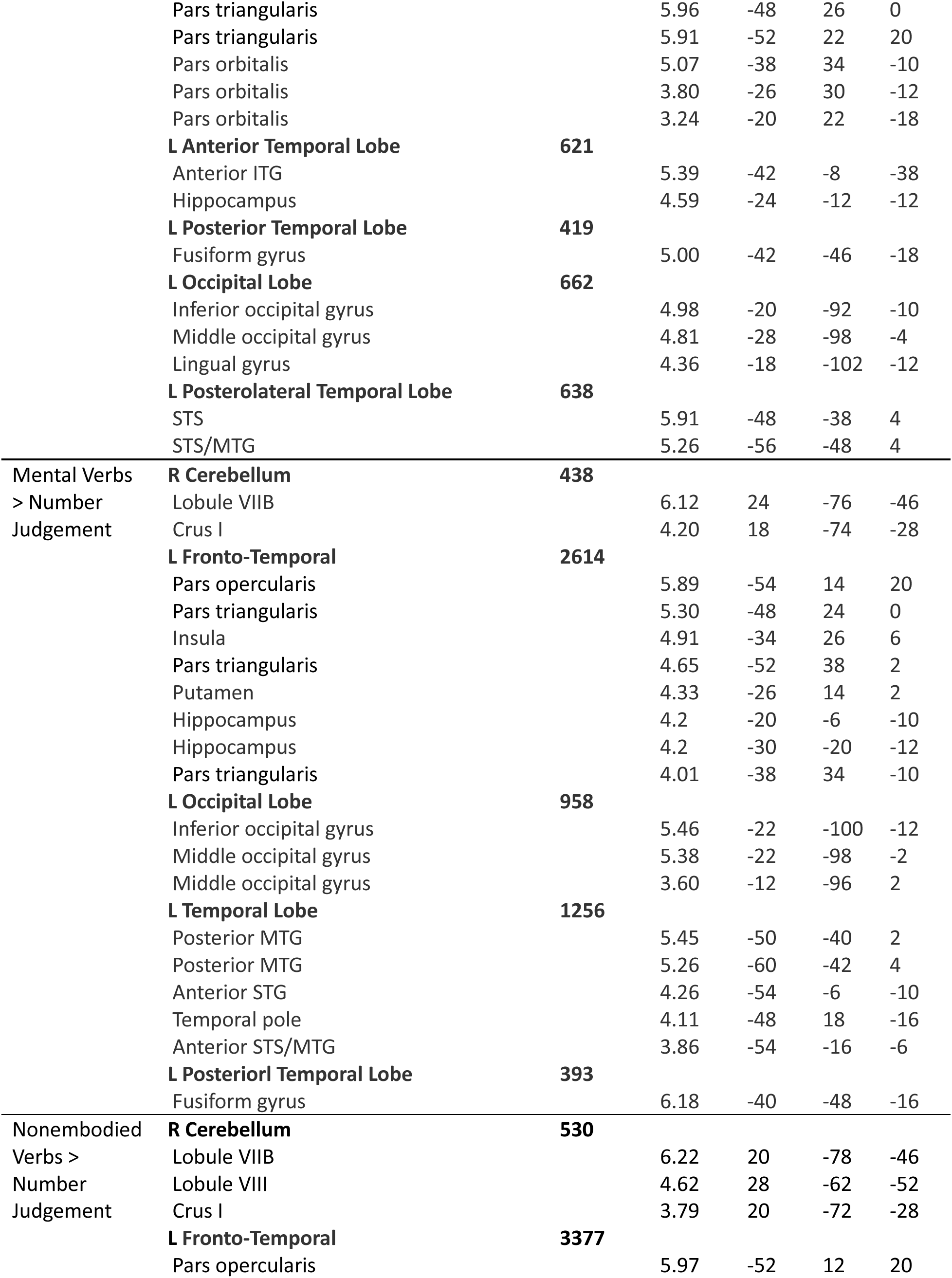
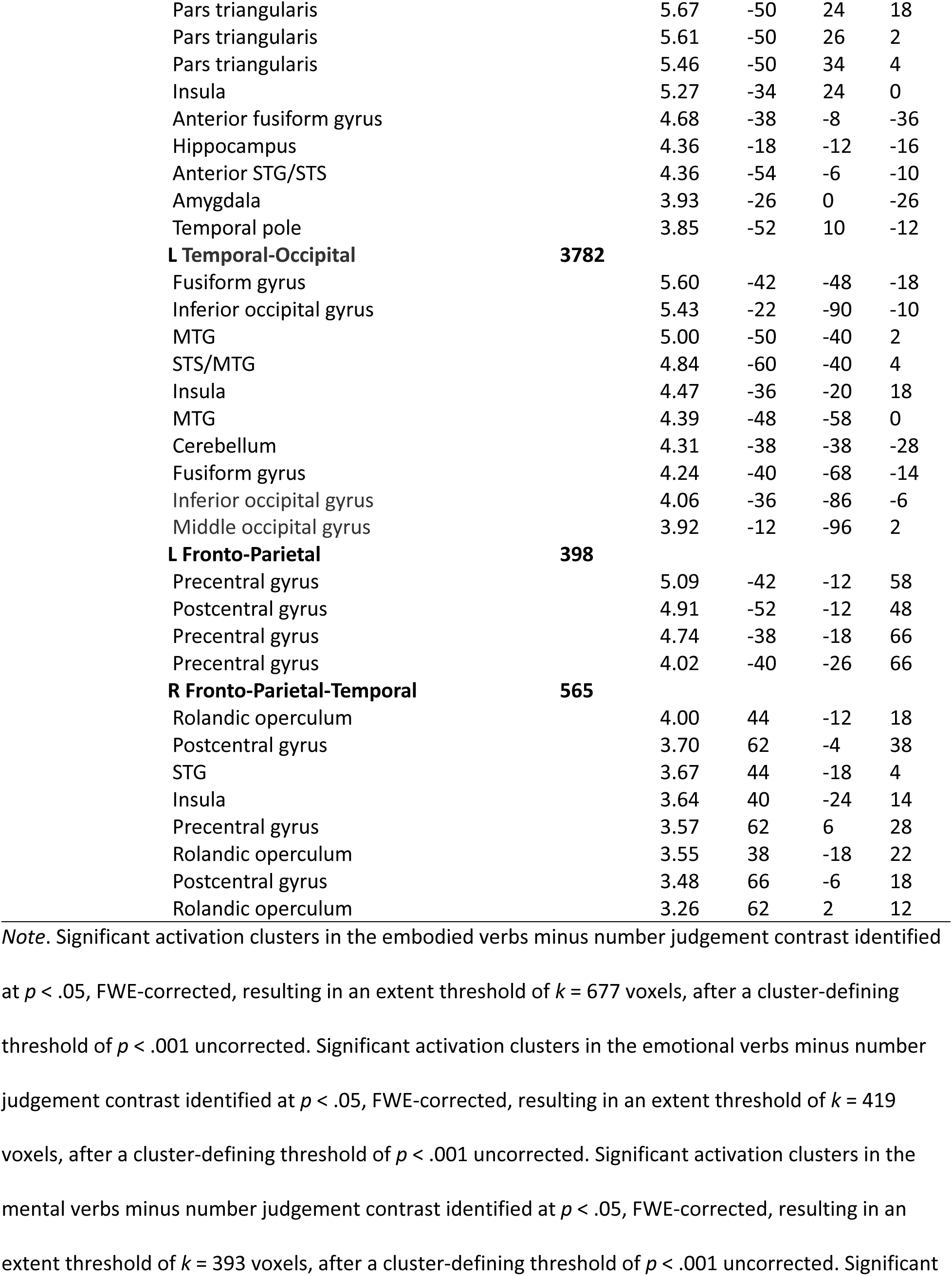
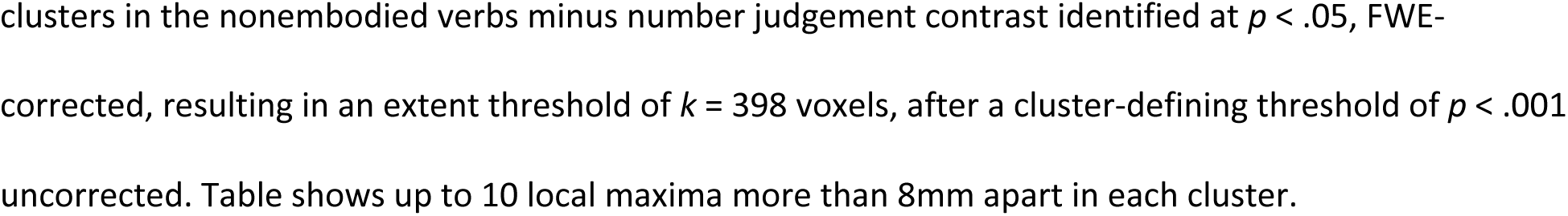
Significant Activation Clusters in the Whole Brain Analyses.

### ROI Analyses

Next, we conducted a series of focused a priori ROI analyses (see Figure 3a for locations and descriptions). The ROIs were defined based on peak activation coordinates from previous literature, and were selected to correspond to the semantic categories of our verb stimuli, as well as semantic processing more generally. In the ventral ATL we observed significant activation for all verb types in contrast to the number judgement task, apart from the mental verbs (Figure 3b) that did not activate the ventral ATL to a significantly greater degree than in the number judgement task. We observed no significant differences in activation when comparing abstract verb types to embodied verbs in ROIs previously associated with emotional, mental, and nonembodied information (Table 9). However, when comparing the embodied verbs to nonembodied verbs activity in the left posterior STS was marginally significant (*p* = .051). This ROI has been associated with processing motion related concepts (Kuhnke et al., 2022).

**Figure 3.**
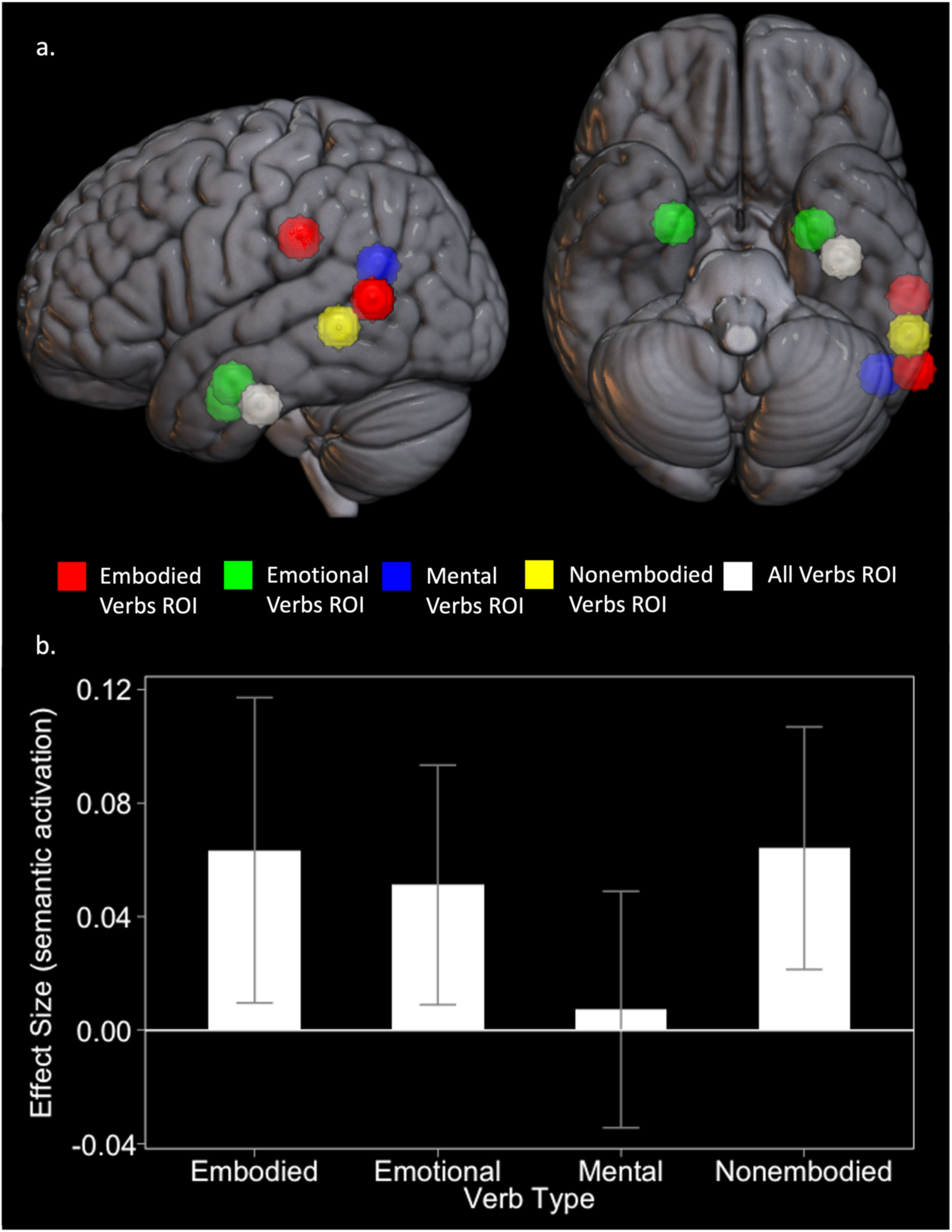
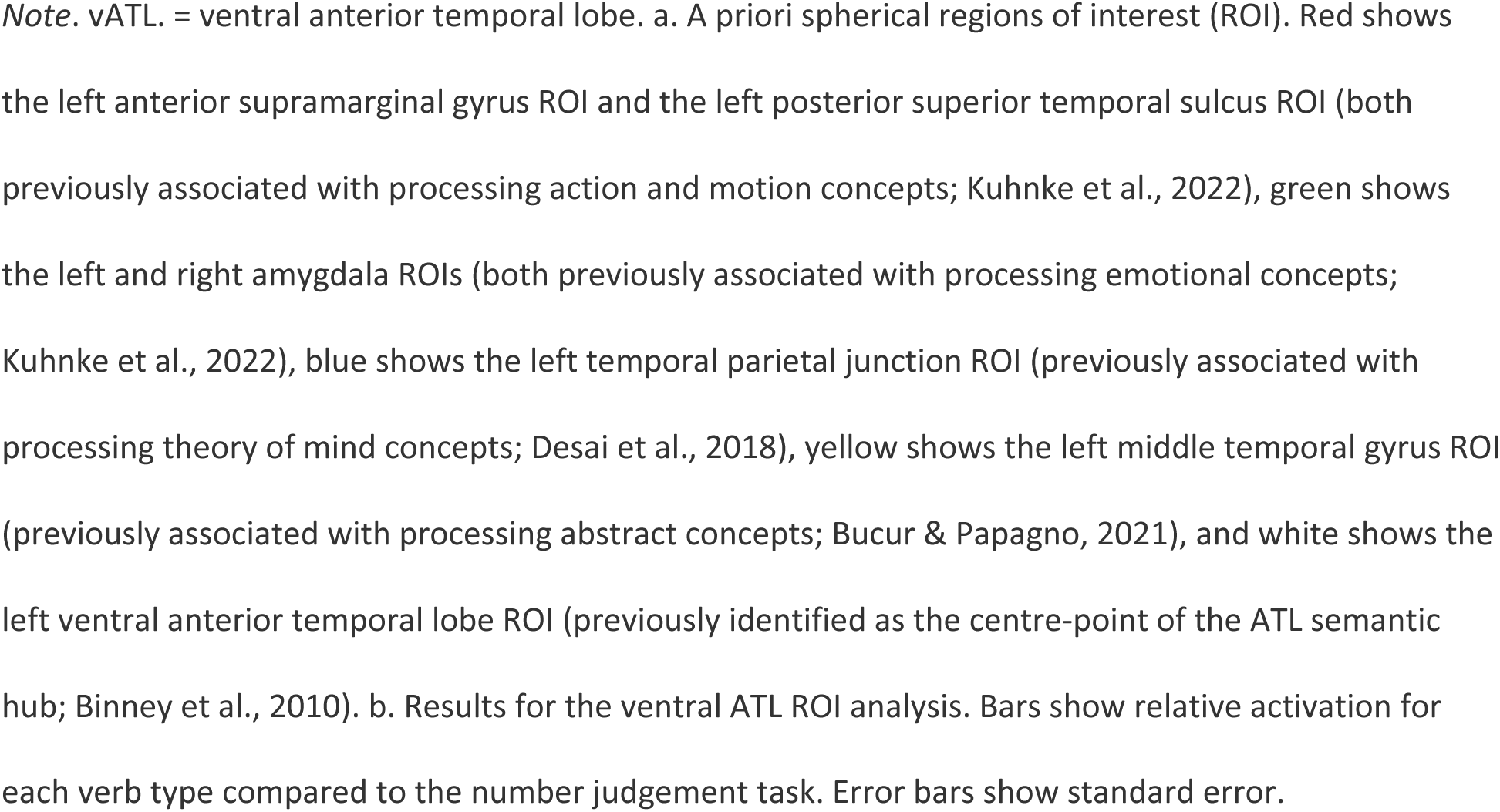
Spherical Region of Interest Locations and Contrast Estimates for the vATL.

**Table 9.** Region of Interest Contrast Results.

| Contrast | ROI | *t* | *p* |
| --- | --- | --- | --- |
| Embodied > Nonembodied | L. anterior supramarginal gyrus | 1.00 | .163 |
|  | L. posterior superior temporal sulcus | 1.70 | .051 |
| Emotional > Embodied | L. amygdala | -3.00 | .997 |
|  | R. amygdala | -2.85 | .996 |
| Mental > Embodied | L. temporal parietal junction | -1.76 | .955 |
| Nonembodied > Embodied | L. middle temporal gyrus | -0.90 | .813 |
| Embodied > Number | L. ventral ATL | 1.94 | .031* |
| Emotional > Number | L. ventral ATL | 2.00 | .028* |
| Mental > Number | L. ventral ATL | 0.29 | .387 |
| Nonembodied > Number | L. ventral ATL | 2.47 | .010* |

*Note.* L = left, R = right, ATL. = anterior temporal lobe.

### Multivariate Analyses

A searchlight MVPA restricted to the ATL revealed clusters sensitive to differences between each abstract verb type and embodied verbs (Figure 4 and Table 10). A group of voxels in the bilateral dorsolateral temporal poles was associated with high classification accuracy when differentiating between mental verbs and embodied verbs. A group of voxels in the left middle temporal gyrus in the posterior portion of the ATL was associated with high classification accuracy when differentiating between emotional and embodied verbs. A similarly located group of voxels in the left middle and inferior temporal gyri in the posterior portion of the ATL was associated with high classification accuracy when differentiating between nonembodied and embodied verbs.

**Figure 4.**
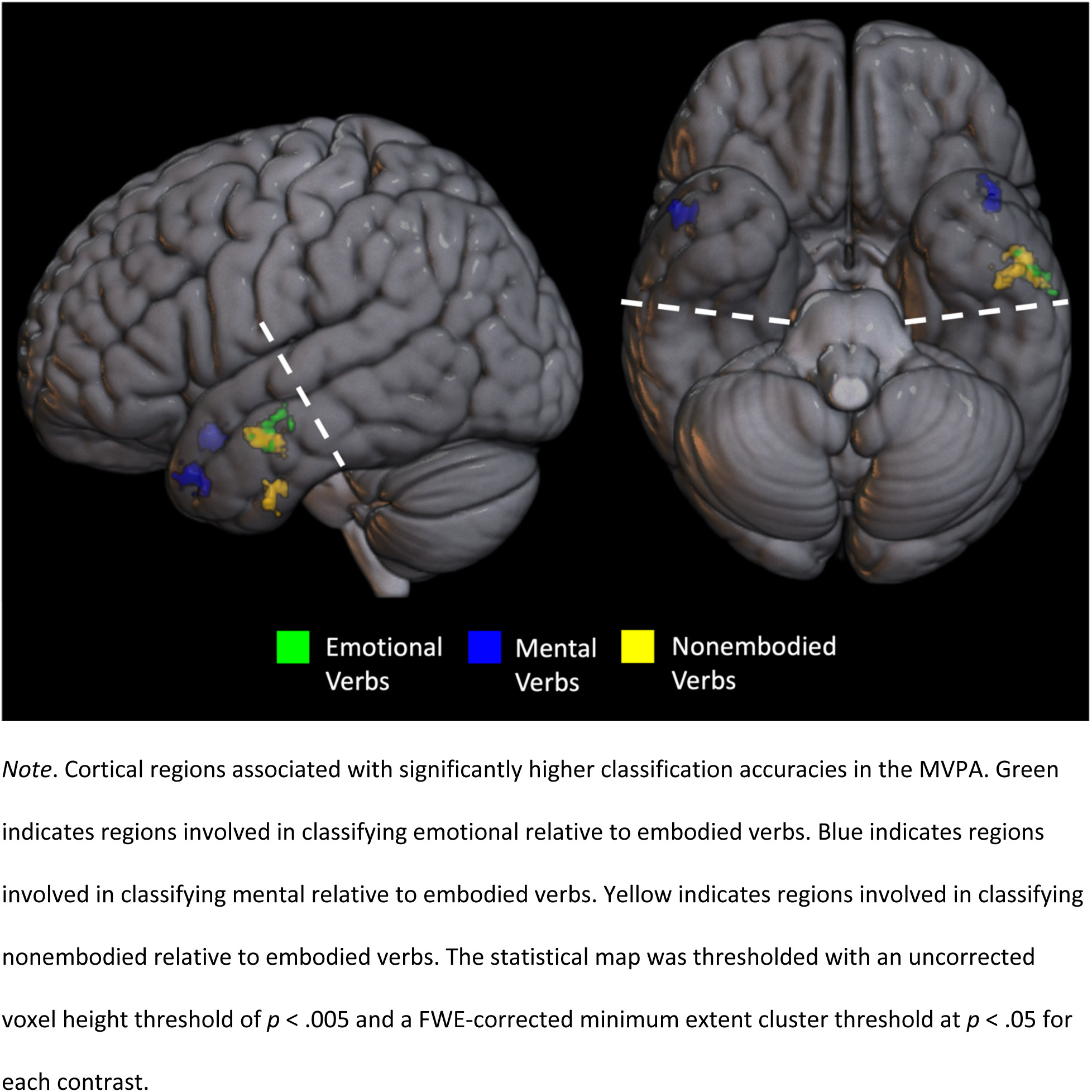
*Note.* Cortical regions associated with significantly higher classification accuracies in the MVPA. Green indicates regions involved in classifying emotional relative to embodied verbs. Blue indicates regions involved in classifying mental relative to embodied verbs. Yellow indicates regions involved in classifying nonembodied relative to embodied verbs. The statistical map was thresholded with an uncorrected voxel height threshold of *p* < .005 and a FWE-corrected minimum extent cluster threshold at *p* < .05 for each contrast.

**Table 10.** Significant Clusters Associated with Classification Accuracy in MVPA.

| Contrast | Cluster name and location of maximum | Cluster extent (voxels) | Peak (Z) | Maxima MNI coordinates (mm) |  |  |
| --- | --- | --- | --- | --- | --- | --- |
|  |  |  |  | x | y | z |
|  | L MTG | 36 |  |  |  |  |
| Emotional ><br>Embodied<br>Verbs |  |  | 3.57 | -54 | -4 | -20 |
|  |  |  | 3.21 | -62 | -8 | -14 |
| Mental ><br>Embodied<br>Verbs | **R Temporal Pole** | **35** |  |  |  |  |
|  |  |  | 3.81 | 54 | 10 | -20 |
|  | **L Temporal Pole** | **31** |  |  |  |  |
|  |  |  | 3.54 | -44 | 12 | -34 |
|  |  |  | 3.31 | -42 | 22 | -32 |
| Nonembodied<br>> Embodied<br>Verbs | **L MTG/ITG** | **40** |  |  |  |  |
|  |  |  | 4.03 | -54 | -4 | -22 |
|  |  |  | 2.96 | -60 | -12 | -26 |
|  | **L ITG** | **32** |  |  |  |  |
|  |  |  | 3.57 | -46 | -12 | -36 |
|  |  |  | 3.07 | -52 | -6 | -42 |

*Note.* Significant activation clusters in emotional relative to embodied verb accuracy identified at $p < .05$ , FWE-corrected, resulting in an extent threshold of $k = 36$ voxels, after a cluster-defining threshold of $p < .005$ uncorrected. Significant activation clusters in mental relative to embodied verb accuracy identified at $p < .05$ , FWE-corrected, resulting in an extent threshold of $k = 31$ voxels, after a cluster-defining threshold of $p < .005$ uncorrected. Significant activation clusters in nonembodied relative to embodied verb accuracy identified at identified at $p < .05$ , FWE-corrected, resulting in an extent threshold of $k = 32$ voxels, after a cluster-defining threshold of $p < .005$ uncorrected. Table shows up to 2 local maxima more than 8mm apart in each cluster.

## Discussion

There is growing consensus that multiple representation theories better account for semantic processing of both concrete and abstract concepts, compared to theories which propose that conceptual representations are solely amodal, or solely grounded in sensorimotor systems. The hub-and-spoke hypothesis is a neurobiological and computational model of how meaning, including that of words, can be derived from sensory, motor, emotional, and linguistic experience (Lambon Ralph et al., 2017); modal cortical and subcortical spoke regions provide entry/exit points for semantic information that is integrated and abstracted into supramodal representations subserved by the ATL hub. Previous direct tests of the hub-and-spoke hypothesis have focused predominantly on concrete nouns, with just a few investigations of abstract concepts. Its potential to account for verbs, and particularly abstract verbs, has not previously been investigated with fMRI. Hence, this was the aim of the present study. The key findings were as follows.

1. The vATL was engaged in semantic processing of all verb types (except mental state verbs),
2. Outside of the ATL, we observed robust activity within other parts of core semantic cognition
3. We observed only modest differences between verb types in cortical activity outside of the core semantic network. Nonembodied abstract verbs (e.g., *reduce*) were associated with activity in the motor and somatosensory cortices. Embodied verbs (e.g., *straddle*) were associated with posterior STG, at least to a greater extent than nonembodied verbs (though these differences representations and provide new insight into the role of the wider ATL region in abstract verb processing. We found only modest evidence for differential activity associated with different verb types across distributed ‘spoke’ regions. However, differential activity observed within the ATL suggests that the kinds of information converging within the ATL are dependent on verb type, and this manifests in graded patterns that are consistent with the graded hub hypothesis.

### The vATL Supramodal Semantic Hub

The whole brain analyses revealed activity in the vATL during the language task when contrasted to the baseline number judgement task, as well as in each of the contrasts between the embodied, emotional, and nonembodied verbs and the baseline task. This confirms a role of the vATL in processing concepts of all types, and not just nouns or concrete concepts (Yi et al., 2007). Previous research employing a range of neuroscience methods has converged on the vATL as a core supramodal semantic hub (Bajada et al., 2019; Binney et al., 2016; Hoffman et al., 2015; Rice et al., 2018) and atrophy of the vATL is associated with profound semantic impairment (Patterson et al., 2007). However, most of this research has focused on concrete nouns, and the role of the vATL in representing abstract concepts and verbs has been less clear.

The neuropsychological data present a conflicted picture, as illustrated by a recent scoping review of abstract vs concrete word processing deficits in patients with vATL atrophy (i.e., semantic dementia); many studies have found that abstract word processing is more preserved than concrete word processing, which suggests the vATL is less important for the representation of abstract concepts, while some studies found the opposite (more preserved concrete word processing) and others found no difference (Mancano & Papagno, 2023). However, these inconsistencies may reflect varying degrees of control for word frequency, individual differences between patients in word familiarity, as well as idiosyncrasies in the distributions of atrophy present in some case studies (Hoffman, 2016). Recent ATL-optimized fMRI studies of healthy adult samples, on the other hand, demonstrate that the vATL is active during processing of abstract as well as concrete concepts (Bajada et al., 2019; Binney et al., 2016; Hoffman et al., 2015; Rice et al., 2018).

Some patient data suggest that the role of the vATL in processing abstract and concrete words could be dependent on syntactic class. For example, Yi and colleagues found that a cohort of semantic dementia patients had greater deficits in concrete motion verb processing relative to abstract cognition verbs, but no such difference for noun processing (Yi et al., 2007). In contrast, Papagno et al. (2009) reported relatively spared processing for both abstract and concrete verbs, and for concrete nouns, but impairments for abstract nouns. Other studies found no differences between concrete and abstract verb processing in semantic dementia (Mancano & Papagno, 2023). The present study was the first to use fMRI to examine the role of the ATL in abstract verb processing. We found an equal, positive response of the vATL to all types of verbs, except for mental state verbs.

Our previous research has found that, during a syntactic classification task like that used here, mental verbs are processed more quickly than other abstract verbs (Muraki et al., 2022). In the scanner, we found trends in this direction, although the only significant difference found was in accuracy and between mental state verbs and emotional verbs. Muraki et al. (2022) also found mental state verbs are more difficult to differentiate from one another in a memory task, relative to other verb types. On this basis, we propose two explanations as to why the vATL was not engaged by abstract mental state verbs. One possibility is that, although our verb stimuli were carefully controlled across multiple dimensions that influence lexical semantic processing, there was an undefined feature of the verbs that was more prominent amongst the mental verbs and made it is easier to classify these words as verbs, on average (e.g., form typicality or discrepancy; de Zubicaray et al., 2023; Sharpe & Marantz, 2017). In turn, that could mean there is less need to engage semantic processes to arrive at a decision. Alternatively, there may be underlying differences in their representation which could facilitate syntactic classification and preclude a requirement for deeper semantic analysis. For example, verbs describing mental processing (e.g., *realize*, *foresee*, *ponder*) might, by virtue of being less differentiated and/or more semantically diverse, trigger rapid retrieval of a general sense of meaning that is sufficient for making the syntactic judgment, but doesn’t necessarily require delving into specific representations subserved by the vATL. Another possible mechanism is that mental state verbs have privileged access to other cognitive systems like the theory of mind network and this facilitates lexical level processing. If either of these accounts were correct, then more demanding semantic tasks (e.g., pairwise relatedness judgments) should evoke vATL activation for mental state verbs.

### The Graded ATL Semantic Hub

The MVPA analyses revealed several clusters of voxels over which activity can be used to classify each abstract verb type, relative to the embodied verbs. This overall pattern is consistent with a graded semantic hub account that characterizes anterior temporal cortex as a unified representational space, but one that exhibits gradients of connectivity that give rise to shifts in semantic function (Bajada et al., 2019; Binney et al., 2012; Rice et al., 2015). At the center of this space lies the ventrolateral ATL, which is engaged by all concept types and semantic information of any kind. Towards the edges, there are gradual shifts in semantic function such that regions on the periphery are relatively more specialized for (i) receiving inputs from particular modalities (e.g., vision or audition), and (ii) encoding certain types of semantic features (for a computational exploration of this general hypothesis, see Plaut, 2002).

We observed that mental state verbs are associated with voxels in the bilateral temporal poles, and this could align with prior studies that attribute a role of the poles in processing social concepts (Balgova et al., 2022, 2024; Binney et al., 2016; Lin et al., 2018; Olson et al., 2007). Specifically, the mental state verbs in the present study are related to cognition, and defined as mental actions or processes of acquiring knowledge (Muraki et al., 2022). Thus, the meaning of many of these verbs could include socially relevant information, such as information about one’s own and others’ mental states, driving involvement of the poles.

In the present study, emotional verbs were associated with an ATL subregion located in the anterior to middle MTG of the left hemisphere. Previous studies have found that emotional semantic information drives bilateral temporal pole activation, in a similar way to social concepts, and it has been suggested that this reflects direct connectivity of the poles with limbic regions (Binney et al., 2012; X. Wang et al., 2019). The region identified here is more posterior and has been associated with processing more general socioemotional qualities of concepts (Hung et al., 2020; X. Wang et al., 2019). It is also more broadly implicated in social cognition, along with the superior temporal sulcus (Deen et al., 2015; Jackson et al., 2023), and this has been linked to the MTG’s role as part of a default mode network (Hung et al., 2020) that supports internally-generated (perceptually-decoupled) cognitive processes, such as mind-wandering, self-referential activities, autobiographical memory, and emotional processing (Raichle, 2015; Satpute & Lindquist, 2019). Satpute and Lindquist (2019) propose that the function of the default mode network in emotional processing is to abstract away from physiological features of emotional experience and encode the multimodal conceptualization of an emotional category.

Finally, nonembodied verbs were associated with a cluster slightly anterior to that for emotional verbs, as well as a more ventromedial cluster of voxels in the ITG. ITG involvement could reflect graded ATL functional sensitivity to visual semantic information (Hoffman et al., 2015). Given that nonembodied verbs refer to actions, states, or relations that occur external to the human body, it is possible that they are associated with more perceptual experience and this drives inputs to the inferior ATL from adjacent visual association regions. All the above possibilities will need to be explored further in future research.

### Differential Engagement of Modality-specific Spokes for Different Categories of Verbs

According to the hub-and-spoke account of semantic representation, the ATL hub shares reciprocal connections with a network of modality-specific regions (e.g., visual, auditory, somatosensory cortices), and coherent conceptual representations arise from the conjoint computations of both the hub and these distributed spokes. Multidimensional theories further posit that engagement of each modal system, during both encoding and retrieval, will vary with the extent to which different types of sensory, motor, emotional, and linguistic information contributes to a representation. fMRI studies and meta-analyses, and a neurostimulation study, support this notion (Conca et al., 2021; Desai et al., 2018; Guo et al., 2013; Kuhnke et al., 2022; Pobric et al., 2010; Reilly et al., 2016). In the present study, we compared categories of concepts that varied in the degree to which sensorimotor, emotional, and other interoceptive experience (i.e., cognition) contributes to their representation (according to ratings studies). Contrary to previous studies, we found only weak evidence of category-specific activation in regions outside of the ATL. This manifested mainly in the extent of activation rather than regional dissociations, with the exception of left precentral gyrus and bilateral postcentral gyrus activity which appeared specific to nonembodied verbs. These findings were, however, only observed in the contrast against the numerical baseline task and were absent from direct comparison between verb types. Overall, our results do not support the hypotheses that emotional verbs are associated with activity in the bilateral amygdalae, mental state verbs are associated with activity in the left temporal parietal junction, or that nonembodied verbs are associated with activity in the left MTG.

Other than these hypotheses being incorrect, there are several possible alternative explanations for these null findings. The first relates, once again, to the nature of the task demands, and the possibility they do not drive the semantic system hard enough to elicit robust differences between categories. Afterall, participants were asked to discriminate verbs from nouns, not between different types of verbs, and this may have limited the depth of semantic analysis required. This would appear inconsistent, however, with the finding that most verb types engaged the vATL. A second possibility is that there is a high degree of heterogeneity amongst concepts within a category, and this drives high variability of engagement both across and within spoke regions which will be lost to signal averaging. The emergence of differential downstream activity across ATL subregions (see above) is not incompatible with this, as it could reflect convergence and summation of inputs from several pathways. Finally, a third possibility is that all abstract verbs are heavily reliant on linguistic information and minimally engage sensorimotor, affective, or interoceptive neural systems. Engagement of language systems in processing abstract concepts was associated with activation of the left inferior frontal gyrus by Hoffman and Bair (2024) in their recent large meta-analysis of neuroimaging studies and with the pars orbitalis of the IFG and left anterior STS by Binder et al. (2009) in their meta-analysis of neuroimaging studies. Here, we observed IFG activation in response to all abstract verb types. Abstract words also tend to occur in more diverse semantic contexts (Hoffman et al., 2013), and require a greater degree of cognitive control to ensure the correct meaning is retrieved (Hoffman et al., 2015; Hoffman & Bair, 2024). Semantic control has also been associated with the IFG. However, these explanations would appear somewhat incompatible with another of our findings. Contrary to our expectations that the nonembodied verbs would be the most abstract, and therefore the most reliant on linguistic representations, they were the only verbs associated with activation of sensorimotor neural areas. One possible interpretation of this is that our nonembodied verbs evoke event representations which engage sensorimotor systems (Grush, 2004; Schütz-Bosbach & Prinz, 2007). For example, many of these stimuli referred to changes of state that could *involve* the human body even if they do not *affect* the body (e.g., *uncork*, *ignite*, *tenderise*). These verbs have low embodiment ratings, but may, nonetheless, include action-related sensorimotor information as part of their meanings.

### Conclusions and Future Directions

To our knowledge, the present study is the first to test the hub-and-spoke account in relation to abstract verb processing. Our findings extend previous research, providing novel evidence that the vATL is engaged in embodied and nonembodied verb processing, adding to the list of concept types that appear to be represented by the vATL. Further, we found evidence to support the graded semantic hub hypothesis, namely differential activity of ATL subregions for different types of verbs. We found little support for strong modality-specificity associated with the verbs, in that we were unable to obtain evidence for differential activity in distributed modality-specific regions. Although the verb stimuli were carefully selected, the categories of abstract verbs used here could fail to adequately capture heterogeneity in the recruitment of modality-specific systems. This may suggest that abstract verbs draw upon multiple, overlapping sources of information and experience. Future research could explore this via representational similarity analyses that correlate patterns of brain activity with a richer and continuous set of information about the dimensions underlying abstract word meaning.

